# Tracing the evolutionary emergence of the temperature sensing prion-like domain in EARLY FLOWERING 3 across the plant kingdom

**DOI:** 10.1101/2023.12.07.570556

**Authors:** Zihao Zhu, Jana Trenner, Marcel Quint

## Abstract

Plants have evolved to anticipate and adjust their growth and development in response to environmental changes. To mitigate the negative influence of global climate change on crop production, understanding the key regulators of plant performance is imperative. *EARLY FLOWERING 3* (*ELF3*) is such a regulator involved in the circadian clock and thermomorphogenesis. *Arabidopsis thaliana* ELF3 contains a prion-like domain (PrLD) that functions as a thermosensor, enabling its liquid-liquid phase separation at high ambient temperatures. To understand the conservation of this function across the plant kingdom, we traced the evolutionary emergence of ELF3 with a focus on PrLD existence. We observed that the presence of the domain within ELF3, mainly contributed by the length of polyglutamine (polyQ) repeats, is most prominent *Brassicales*. By analyzing 319 natural *Arabidopsis thaliana* accessions, we detected a wide range of polyQ length variation in ELF3. However, it is only weakly associated with geographic origin, climate conditions and classic temperature-responsive phenotypes. Based on available prediction tools and limited experimental evidence, we conclude that although the emergence of PrLD is not likely to be a key driver of environmental adaptation, it adds an extra layer to ELF3’s role in thermomorphogenesis.

## Introduction

Plants, like other organisms on Earth, experience both predictable and unpredictable environmental changes. While the regular light/dark and warm/cool cycles can be anticipated by the plants’ internal circadian clock, unpredictable global climate change is demanding their ability to acclimate for evolutionary adaptation. Understanding the key players involved in these processes will help to increase the fitness in crops and mitigate the negative influence of climate change.

As plants are more frequently encountering predictable environmental changes, circadian anticipation is a fundamental attribute contributing to plant performance. The plant circadian clock is composed of multiple interconnected transcriptional-translational feedback loops (Huang and Nusinow, 2016; Nohales and Kay, 2016). These loops can be classified in a time-of-day dependent manner based on the phase of involved clock components. The morning loop contains CIRCADIAN CLOCK ASSOCIATED 1 (CCA1) and LATE ELONGATED HYPOCOTYL (LHY) which positively regulate the expression of *PSEUDO-RESPONSE REGULATOR 9* (*PRR9*) in the morning and *PRR7* in the afternoon (Farré et al., 2005; Nakamichi et al., 2010), while repressing two additional afternoon-phased genes, *PRR5* and *GIGANTEA* (*GI*) (Lu et al., 2012; Kamioka et al., 2016). PRR9, PRR7, and PRR5 later repress the expression of *CCA1* and *LHY*, allowing the induction of evening-phased genes (Nakamichi et al., 2010; Adams et al., 2015). At dusk, accumulation of TIMING OF CAB EXPRESSION 1 (TOC1) suppresses *GI*, which subsequently triggers the activation of *TOC1* (Kim et al., 2007). In addition, three evening-phased proteins EARLY FLOWERING 3 (ELF3), EARLY FLOWERING 4 (ELF4), and LUX ARRYTHMO (LUX) accumulate and form a protein complex known as the evening complex (EC) (Hsu et al., 2013). The EC directly represses the transcription of *PRR9*, *PRR7*, and *GI*, resulting in the accumulation of CCA1 and LHY before dawn (Nusinow et al., 2011; Herrero et al., 2012; Ezer et al., 2017). With this endogenous network, external cues (known as *Zeitgeber*) can be used as timing input to precisely generate internal biological rhythms. However, not all circadian clock components can serve as a *Zeitnehmer* with the ability to receive the timing information from the *Zeitgeber*. Recent studies identified *ELF3* and *GI* as essential *Zeitnehmers* for clock entrainment to photoperiod signals (Anwer et al., 2020), whereas *ELF3* alone can function as a temperature *Zeitnehmer*, sensing warm/cool cycles (Zhu et al., 2022).

While the circadian clock confers the ability to handle daily environmental fluctuations, plants still encounter challenges from climate change, for instance the rise in ambient temperatures. Plants can acclimate rapidly to elevated temperatures through various adjustments in their morphology and development, collectively known as thermomorphogenesis (Delker et al., 2014; Quint et al., 2016). In *Arabidopsis thaliana* seedlings, these adjustments include elongated hypocotyls and leaf upward bending, which are known to improve cooling capacity (van Zanten et al., 2009; Crawford et al., 2012). As a central regulator of thermomorphogenesis signaling, PHYTOCHROME INTERACTING FACTOR 4 (PIF4) accumulates at warm temperatures and activates auxin biosynthesis genes, promoting cell elongation in petioles and hypocotyls, leaf hyponasty, as well as flowering (Franklin et al., 2011; Kumar et al., 2012; Park et al., 2019). In this thermomorphogenesis pathway, the function of PIF4 is gated by temperature sensing systems and the circadian clock. The photoreceptor phytochrome B (phyB) was the first identified plant temperature sensor (Jung et al., 2016; Legris et al., 2016). Warm temperature accelerates the dark/thermal reversion of phyB from its active Pfr form to its inactive Pr form (Delker et al., 2017). The active Pfr form of phyB mediates PIF4 degradation by stabilizing ELF3 (Nieto et al., 2015). ELF3 contains a prion-like domain (PrLD) which also functions as a thermosensor, enabling liquid-liquid phase separation (LLPS) of ELF3 from its dilute phase into liquid droplets (dense phase) at high temperatures (Jung et al., 2020; Hutin et al., 2023). The dense phase aggregation of ELF3 coordinates with its restricted mobilization to the nucleus (Ronald et al., 2021; Ronald et al., 2022), which potentially relieves the direct interaction with PIF4 (Nieto et al., 2015) and the transcriptional repression of *PIF4* by the EC (Box et al., 2015; Raschke et al., 2015). With its multiple functions connecting temperature sensing, circadian clock, and thermomorphogenesis, *ELF3* has been described as a key plasticity gene contributing to plant acclimation (Blackman, 2017; Laitinen and Nikoloski, 2019).

Expanding knowledge generated from *Arabidopsis thaliana* to crops is necessary to achieve crop-level adaptations and yield stability under global climate change (Challinor et al., 2014). Natural variation or loss-of-function in *ELF3* generally affects circadian clock regulated photoperiodic flowering in various crop species, including rice (Matsubara et al., 2012; Saito et al., 2012; Andrade et al., 2022), barley (Faure et al., 2012; Zakhrabekova et al., 2012; Zahn et al., 2023), wheat (Alvarez et al., 2016; Alvarez et al., 2023; Mizuno et al., 2023; Wittern et al., 2023), soybean (Lu et al., 2017; Bu et al., 2021; Fang et al., 2021), and chickpea (Ridge et al., 2017). This allows the cultivation of crops under altered photoperiods, therefore important for crop domestication and spatial distribution. Besides the clock function, *ELF3* is involved in barley morphological and developmental acclimations to high ambient temperatures (Ford et al., 2016; Ejaz and von Korff, 2017; Zhu et al., 2023), suggesting conserved roles in temperature responsiveness. However, unlike *Arabidopsis thaliana*, the monocot grass *Brachypodium distachyon* does not have a temperature-responsive PrLD in ELF3 (Jung et al., 2020). Therefore, the conserved functions of *ELF3* in relation to temperature sensing remain unclear, particularly in monocots.

In *Arabidopsis thaliana* ELF3, the temperature sensing PrLD harbors natural variation in the length of a polyglutamine (polyQ) stretch caused by expanded cytosine-adenine-adenine (CAA) repeats (Undurraga et al., 2012). In a manner similar to how ELF3 aggregates in response to high temperatures (Jung et al., 2020), it has been observed that in humans, polyQ-extended proteins tend to aggregate in degenerated neurons, leading to the development of polyQ diseases (Fan et al., 2014). This consistency suggests that polyQ determines the thermosensing function of *Arabidopsis thaliana* ELF3-PrLD. However, the potential effects and evolutionary significance of ELF3-polyQ variation in plants are still unknown, even in the model *Arabidopsis thaliana*. In this study, we attempt to shed some light on the evolutionary trajectory of *ELF3*. To assess this in a systematic manner, we traced the evolutionary emergence of *ELF3* across the plant kingdom, with a focus on PrLD existence. Based on 319 *Arabidopsis thaliana* accessions, we sought to examine the correlation between ELF3-polyQ variation and geographic origins, local environments, as well as temperature-responsive phenotypes. Lack of reliable phenotype-polyQ correlations together with the agglomerated presence of the ELF3-PrLD within the *Brassicales* order suggest that it cannot be considered as a general evolutionary adaptation to diverse environments.

## Results

### Evolutionary emergence of *ELF3* and its prion-like domain

In *Arabidopsis thaliana*, the major functions of ELF3 in circadian clock regulation require the involvement of its EC partners ELF4 and LUX (Nusinow et al., 2011; Ezer et al., 2017). Regarding the emergence of the EC, previous studies revealed a homologue of *ELF3* in charophyte *Klebsormidium nitens*, whereas potential homologues of *ELF4* and *LUX* were identified even in the chlorophytes like *Chlamydomonas reinhardtii* (Linde et al., 2017). In order to obtain a general picture about the evolution of *ELF3* and its duplicate *ESSENCE OF ELF3 CONSENSUS* (*EEC)* (Liu et al., 2001), whose function remains unknown, across the plant kingdom, we first determined the copy number of the EC components *ELF3*, *ELF4*, and *LUX*, as well as *EEC* in 42 plant genomes ranging from unicellular green algae to flowering plants (Table 1). An *ELF3* homologue was identified in the charophyte *Chara braunii*, confirming the origin of *ELF3* in Charophyta (Linde et al., 2017). Interestingly, in contrast to the identification of the EC components back to Charophyta, *EEC* homologues emerged later and are restricted to eudicots (Table 1), suggesting that a duplication event of *ELF3* in the last common ancestor of the eudicots gave rise to *EEC* in this lineage.

**Table 1.**
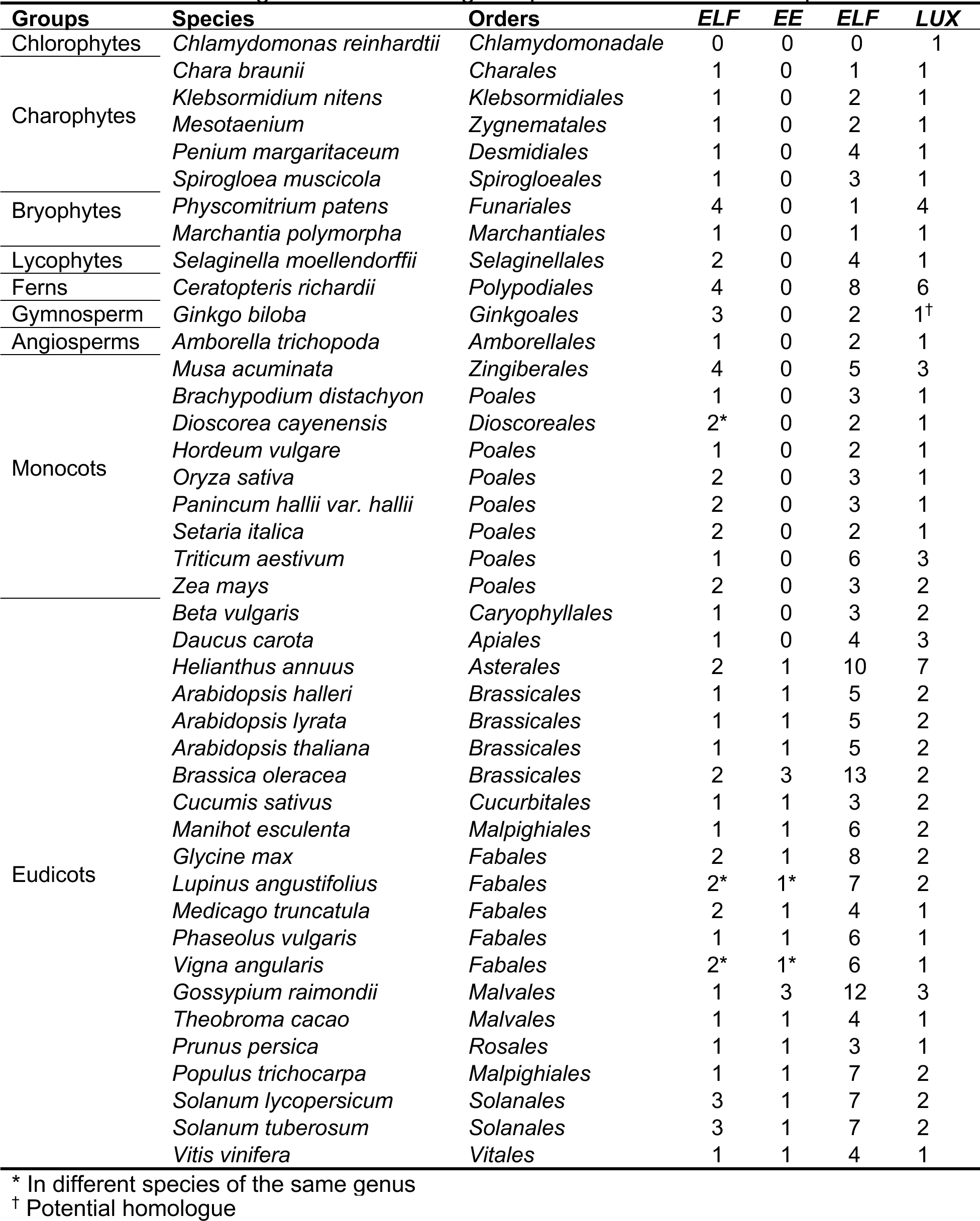
Gene homologues of the evening complex and *EEC* in various plant.

To trace the evolution and divergence of *ELF3* and *EEC* in more detail, the protein homologues of ELF3 and EEC were identified from 274 plant genomes (Supplemental Table S1) and their phylogenetic relationships were reconstructed. The sequences from similar angiosperm groups (e.g., basal angiosperms, monocots, eudicots, and core eudicots) mostly clustered together in the phylogenetic tree (Fig. 1; Supplemental Fig. S1). As expected, ELF3 and EEC were separated into two different clades, with EEC being restricted to core eudicots. In orders such as *Buxales*, *Trochodendrales*, *Proteales*, and *Ranunculales*, which are eudicots but not core eudicots, only ELF3 homologues were detected, positioned in a clade with ELF3 from basal angiosperms and monocots (Supplemental Fig. S1). Interestingly, this clade is more closely related to the EEC clade than to the ELF3 clade from core eudicots. To understand sequence features that distinguish ELF3 and EEC, we next selected 32 species (8 *Poales* and 24 core eudicots that have both ELF3 and EEC homologues, including 7 *Brassicales* species) for sequence alignment. As reported previously (Liu et al., 2001), four highly conserved regions (I-IV) were detected within ELF3 and EEC in these species (Fig. 2). Meanwhile, *Poales* (monocots) ELF3 contained a unique region (VII) and shared a conserved region (VI) with core eudicots EEC in the amino-terminal, potentially separating them from the core eudicots ELF3. Notably, both EEC and ELF3 from *Brassicales* have several unique features (regions V, VI, VIII, IX, and X) compared to the other core eudicots (Fig. 2), in line with the inspection of the branch lengths of *Brassicales* ELF3 and the next closely related core eudicots in the phylogenetic tree (Supplemental Fig. S1). These findings indicate that the *Brassicales* ELF3 sequences are only recently specified and are all rather closely related to each other.

**Fig. 1.**
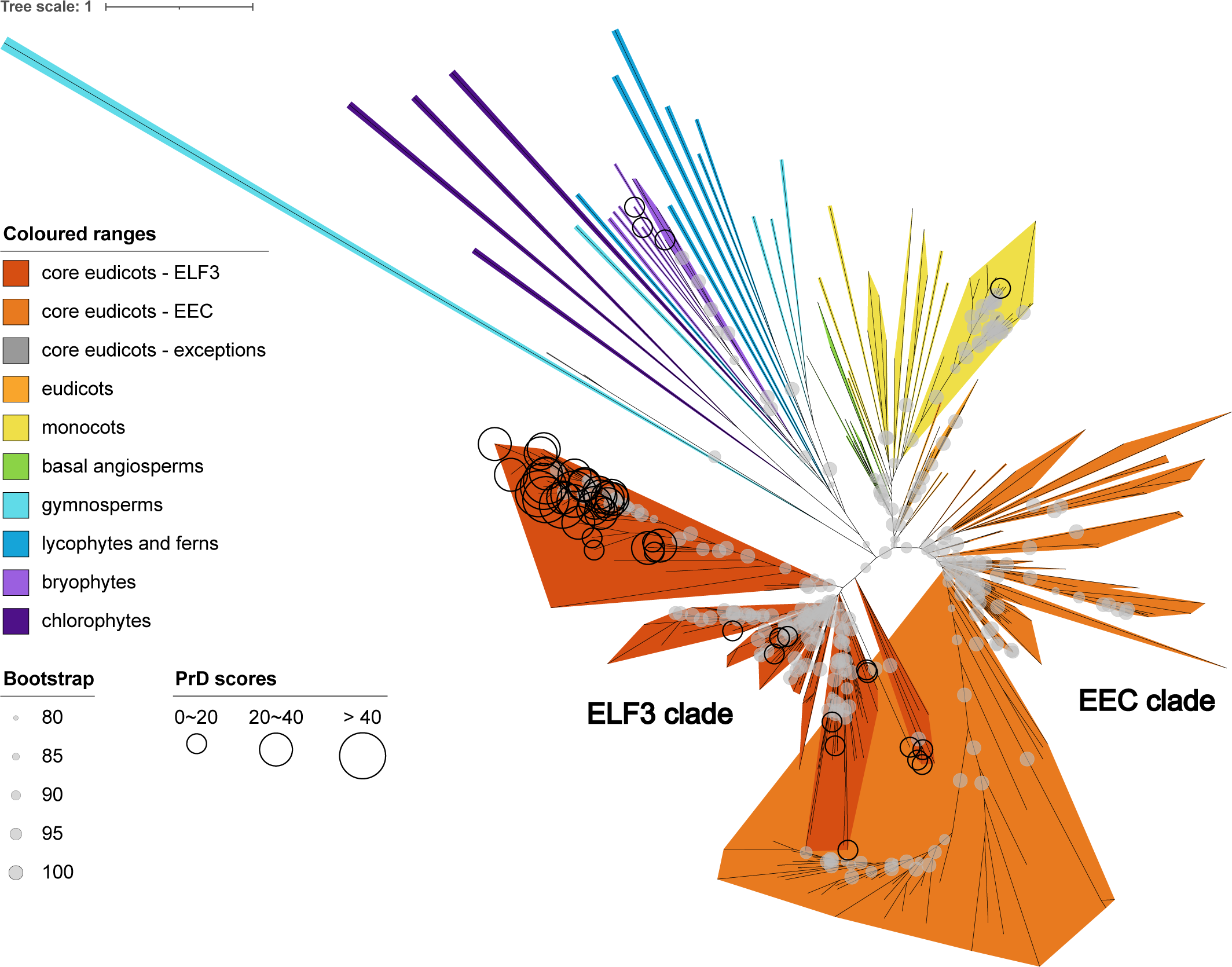
Phylogeny of ELF3 and EEC across the plant kingdom. Phylogenetic tree was constructed with the full-length amino acid sequences obtained from 274 plant genomes, using maximum likelihood IQ-Tree JTT+F+R10 model with 10,000 replications of ultrafast bootstrap (bootstrap values >= 80 are shown as grey circles). ELF3 and EEC clades of core eudicots are marked based on the position of *Arabidopsis thaliana* ELF3 and EEC, respectively. Coloured ranges are based on species group and clade. Sequences with positive PrD scores (PLAAC derived) are marked with black circles with three threshold groups. The center of the circle is placed at the end of the corresponding branch. The number of ‘equal-daylight’ algorithm iteration was set to 1 to increase branch visibility. Full rooted phylogenetic tree with labels is shown in Supplemental Fig. S1.

**Fig. 2.**
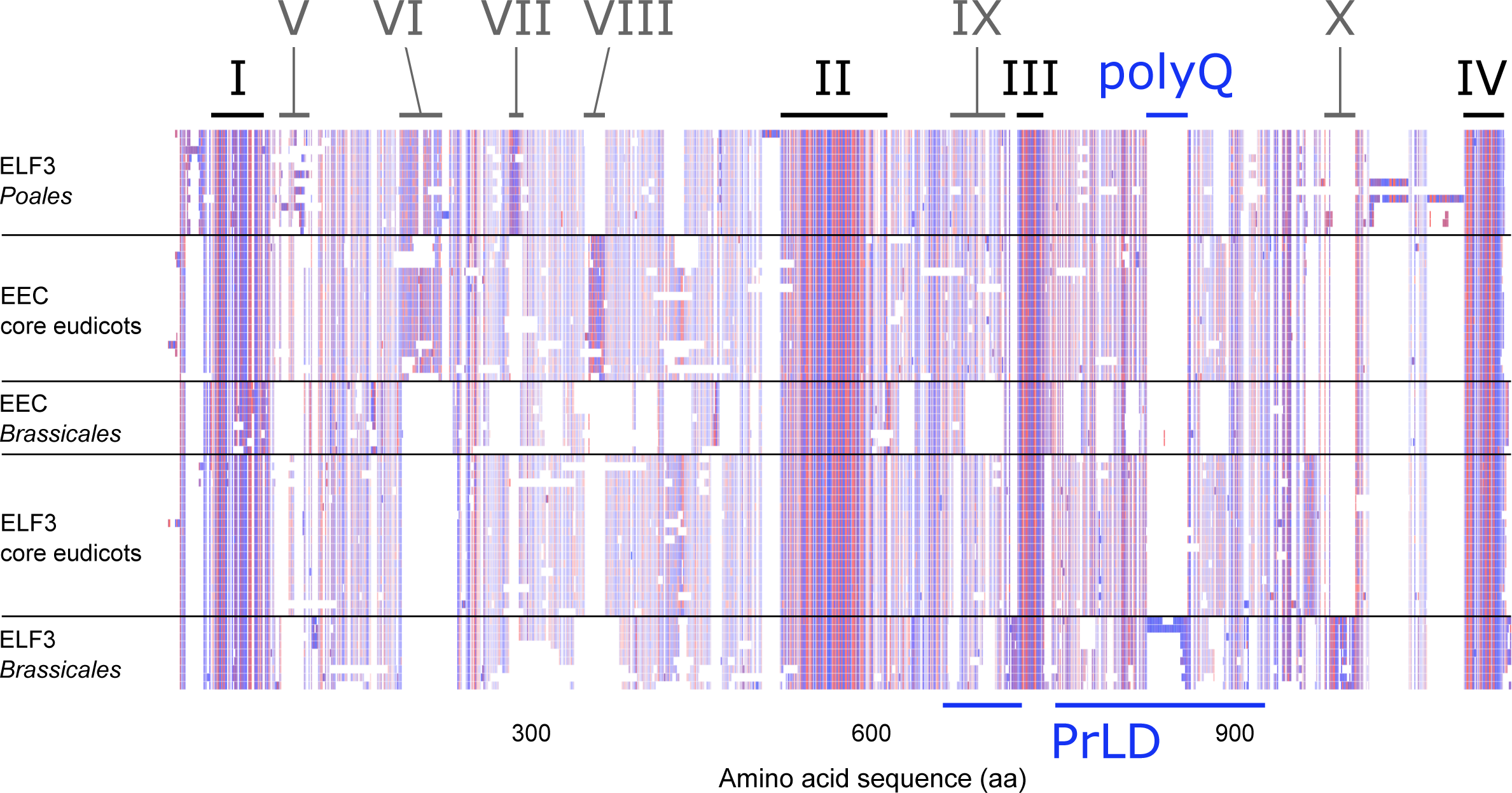
Conserved and distinct features of ELF3 and EEC. Multiple amino acid sequence alignment of ELF3/EEC homologues in eight monocots: from top *Oryza sativa* (2), *Paspalum vaginatum* (2), *Setaria viridis* (2), *Zea mays* (2), *Sorghum bicolor* (2), *Triticum aestivum*, *Hordeum vulgare*, and *Brachypodium distachyon*; 17 core eudicots: *Cicer arietinum* (2 ELF3), *Medicago truncatula* (2 ELF3), *Glycine max* (2 ELF3), *Phaseolus vulgaris*, *Alnus serrulata*, *Carya illinoinensis*, *Castanea pumila*, *Manihot esculenta*, *Populus trichocarpa*, *Salix purpurea*, *Citrus clementina*, *Toxicodendron radicans*, *Bixa orellana*, *Gossypium barbadense* (2 EEC), *Fouquieria macdougalii*, *Cucumis sativus*, and *Eucalyptus grandis*; seven *Brassicales* species: *Alyssum linifolium*, *Capsella rubella*, *Arabidopsis thaliana*, *Malcolmia maritima*, *Eutrema salsugineum*, *Crambe hispanica* (2 EEC and 2 ELF3), and *Brassica oleracea* (2 EEC and 2 ELF3). Hydrophobicity is used for amino acids colour scheme and the colour intensity is based on the sequence conservation. Regions I-IV represent four highly conserved regions between ELF3 and EEC, whereas regions V-X represent those with special features between ELF3 and EEC, and/or between species groups. The PrLD regions are based on the PLAAC analysis of *Arabidopsis thaliana* ELF3 (Supplemental Fig. S3), including a polyQ stretch.

*Arabidopsis thaliana* ELF3 is known to harbor a prion-like domain (PrLD) which is required for phase separation of ELF3 in response to temperature changes (Jung et al., 2020; Hutin et al., 2023). The study of mammal and especially yeast prions resulted in a various prediction tools for PrLDs that are based on two different models. The compositional model searches for a large number of weak interactions distributed along the PrLD with a biased amino acid composition (Toombs et al., 2012; Gil-Garcia et al., 2021). Of importance for the so-called soft-amyloid model is the presence of stretches of mild amyloid potential, embedded within an intrinsically disordered protein region (IDPR) (Sabate et al., 2015; Gil-Garcia et al., 2021). As IDPRs are described to be drivers of biological liquid-liquid phase separation (LLPS) (Uversky, 2019), we first analyzed the sequence composition of the identified ELF3/EEC sequences from 274 plant genomes using localCIDER (Holehouse et al., 2017). All ELF3/EEC sequences were assigned to the class of weak polyampholytes and polyelectrolytes, with the fraction of charged residues (FCR) and the net charge per residue (NCPR) below 0.25, representing hallmarks of intrinsically disordered proteins (Supplemental Fig. S2A). To understand whether the PrLD is conserved in identified ELF3/EEC homologues, we next performed a Prion-Like Amino Acid Composition (PLAAC) search on all sequences and obtained scores (PrD score and Log-likelihood ratio, LLR) indicating the probability of the presence of prion subsequences (Lancaster et al., 2014). Compared to the PrD score, LLR does not impose a hard cut-off. For instance, the PrLD of *Arabidopsis thaliana* ELF3 exhibited an identical PrD score and LLR of 31.53, containing two subsequence regions (Fig. 1; Supplemental Figs. S1, S3). When considering the hard cutoff (PrD score), the PrLD prediction identified ELF3 sequences mostly from core eudicots (Fig. 1; Supplemental Fig. S1). In addition, ELF3 homologues in the bryophytes *Physcomitrium patens* and *Sphagnum fallax*, as well as the monocot *Sorghum bicolor* were predicted to have a PrLD with a relatively low but positive PrD score. Nevertheless, with exception of *Sorghum bicolor* but consistent with a previous report on *Brachypodium distachyon* (Jung et al., 2020), monocots generally lack such a domain in their ELF3. High PrD scores were detected almost exclusively in *Brassicales* ELF3, with several species (*Capsella grandiflora*: 59.89, *Arabidopsis lyrata*: 58.27, *Capsella rubella*: 57.52, *Alyssum linifolium*: 49.96, *Arabidopsis halleri*: 46.36, *Descurainia sophioides*: 45.54, *Brassica rapa*: 32.62, and *Isatis tinctoria*: 32.09) displaying an even higher score than *Arabidopsis thaliana*, suggesting potentially conserved temperature sensing functions of PrLDs across *Brassicales*. Moreover, despite four highly conserved regions between ELF3 and EEC, the third conserved region (III) was situated in the gap of the predicted PrLD (Fig. 2; Supplemental Fig. S3) and the polyQ stretch diverged between *Brassicales* ELF3 and all the other sequences (Fig. 2). As a result, none of the sequences in the EEC clade was predicted to have a PrLD (Fig. 1).

To complement the composition-based PrLD prediction of PLAAC, we also analyzed the ELF3/EEC sequences using the PrionW webserver (Zambrano et al., 2015). PrionW employs a combination of compositional and soft-amyloid model for the prediction of PrLDs. The initial approach using the default settings for Q+N richness of 25% and a pWALTZ cut-off of 73 resulted in only three hits: ELF3 of *Isatis tinctoria* (*Brassicaceae*), *Populus deltoides* and *Populus trichocarpa* (*Salicaceae*). In order to include *Arabidopsis thaliana* ELF3, with its experimentally proven PrLD function known to mediate LLPS (Jung et al., 2020; Hutin et al., 2023), we relaxed the PrionW analysis parameters to a Q+N richness of 18% and a pWALTZ cut-off of 68. The relaxed approach resulted in a set of 54 sequences with predicted PrLD, including 19 *Brassicales* sequences. Fourteen of these overlapped with the PLAAC set of 52 sequences, including 33 *Brassicales* sequences (Supplemental Fig. S2B). In contrast to the PLAAC results, PrionW hits can also be found in the EEC clade (Supplemental Fig. S2C). These combined data using different PrLD prediction models show isolated cases of PrLD emergence in selected species, but an expansion of this domain is restricted to ELF3 homologues across *Brassicales*.

### Polyglutamine repeats contribute to the PrLD of *Brassicales* ELF3

As the prediction of PrLD was mainly restricted to *Brassicales* ELF3, we investigated whether the potential PrLDs of these species are conserved at the sequence level. We constructed a phylogenetic tree with *Brassicales* ELF3 only, separating different families (Fig. 3). As main features of PrLD or prion proteins (Harrison and Gerstein, 2003), we observed a considerable proportion of asparagine (N) and glutamine (Q) in ELF3-PrLD regions based on the sequence alignment (Fig. 3). As previously reported (Undurraga et al., 2012), *Arabidopsis thaliana* ELF3 contained a polyglutamine (polyQ) stretch (with over seven consecutive Qs) in its PrLD. Although such a polyQ stretch is specific to *Brassicaceae* ELF3 and absent from other *Brassicales*, its length correlated positively with the PrLD score (Supplemental Fig. S4). For example, *Capsella grandiflora* with the highest PrD score (59.885) also displayed the longest polyQ stretch (33Q, including four histidine gaps) (Figs. 1, 3; Supplemental Fig. S1). In contrast, the number of asparagines was less variable in the PrLD and did not correlate with the PrD score (Supplemental Fig. S4). Hence, although positive PrD scores were also detected in *Brassicales* families *Cleomaceae* and *Salvadoraceae*, the ELF3-PrLD characteristics as measured by the prediction tools used in this study are mainly contributed by the length of polyQ observed in the family *Brassicaceae*. It is important to note here that the *Arabidopsis thaliana* accession Col-0 used in this phylogenetic tree has 7Q in its polyQ stretch, which is enough for the temperature sensing PrLD function (Jung et al., 2020). Provided that the polyQ stretch contributes to the temperature sensing function (Jung et al., 2020), other *Brassicaceaes* with longer polyQ stretches are likewise expected to display temperature-responsive phase separation function conserved in their respective ELF3 proteins.

**Fig. 3.**
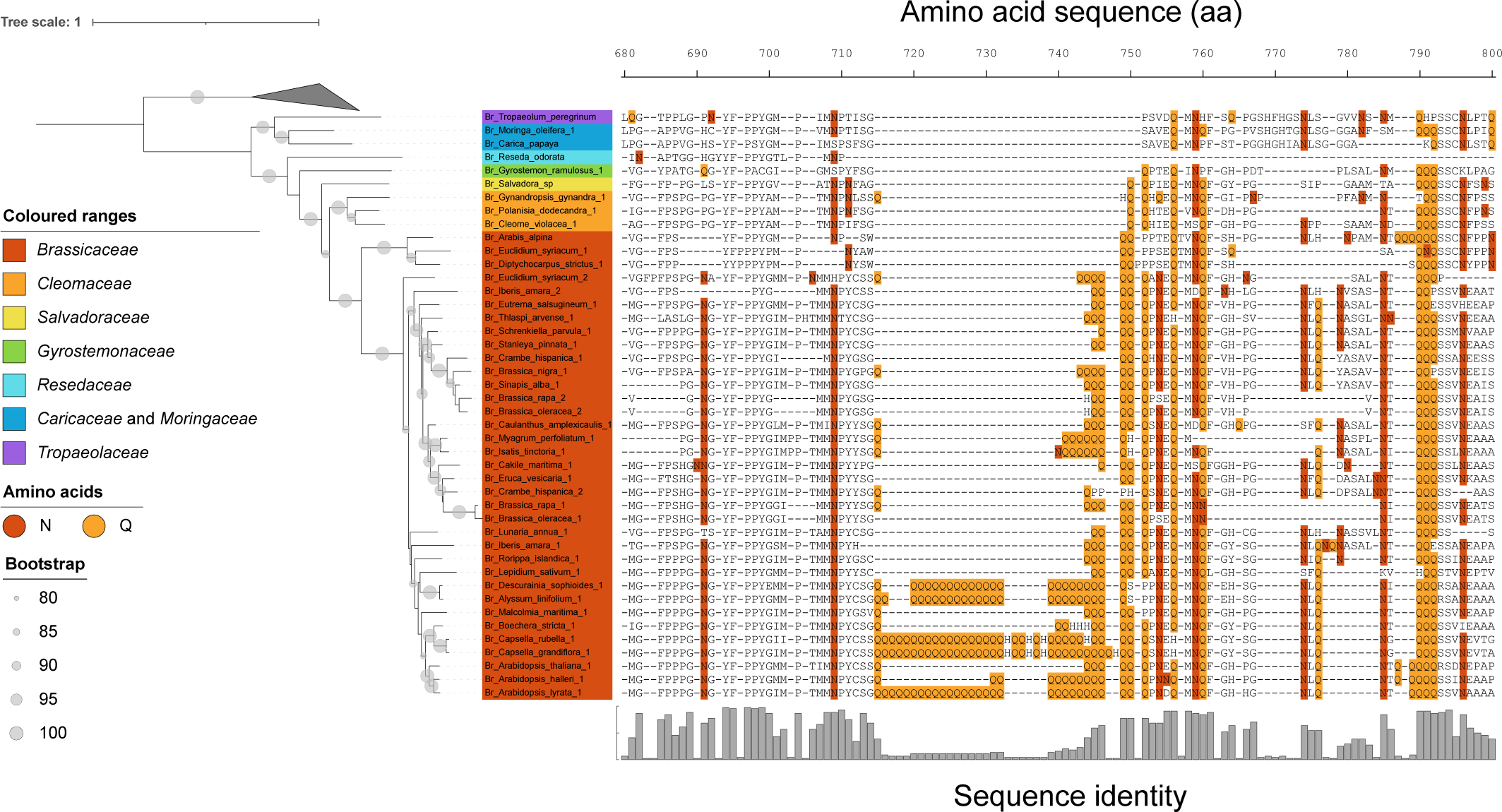
PolyQ stretch in *Brassicales* ELF3. Phylogenetic tree was constructed with the full-length amino acid sequences (44 *Brassicales* and 8 *Poales* species) using maximum likelihood IQ-Tree JTT+R3 model with 10,000 replications of ultrafast bootstrap (bootstrap values >= 80 are shown as grey circles). The *Poales* species were used for rooting and collapsed. The labels are coloured according to the species family. The multiple sequence alignment represents part of the PrLD including the polyQ stretch, with a frequency plot indicating sequence identity. Amino acids asparagine (N) and glutamine (Q) are coloured within the alignment.

### Evolution of Arabidopsis thaliana ELF3 and polyQ

Although data are lacking from most *Brassicaceaes*, natural variation of ELF3-polyQ length has been investigated in several collections of *Arabidopsis thaliana* accessions (Tajima et al., 2007; Undurraga et al., 2012). Likewise, the 1001 Genomes (Alonso-Blanco et al., 2016) provide polymorphism information in *ELF3*, but the polyQ length cannot be identified due to unknown nucleotides in the region, probably caused by common problems of short-read sequencing approaches in highly repetitive regions. Therefore, we dideoxy-sequenced the corresponding region in an additional 204 accessions obtained from the 1001 Genomes collection and corrected their *ELF3* sequences accordingly. As a result, together with previously reported data (Tajima et al., 2007; Undurraga et al., 2012), corrected *ELF3* sequence information was available for further analyses for a total of 319 *Arabidopsis thaliana* natural accessions (Fig. 4A). Among these accessions, ELF3-polyQ length displayed a nearly normal distribution with 16Q being the most frequent, although 15Q and 17Q were rather rare (Fig. 4B). The polyQ length ranged from 7Q to 29Q with a slightly skewed distribution towards <16Q. These data suggest that PrLDs are conserved across *Arabidopsis thaliana* accessions, as its function was originally described using accession Col-0 with the shortest 7Q (Jung et al., 2020).

**Fig. 4.**
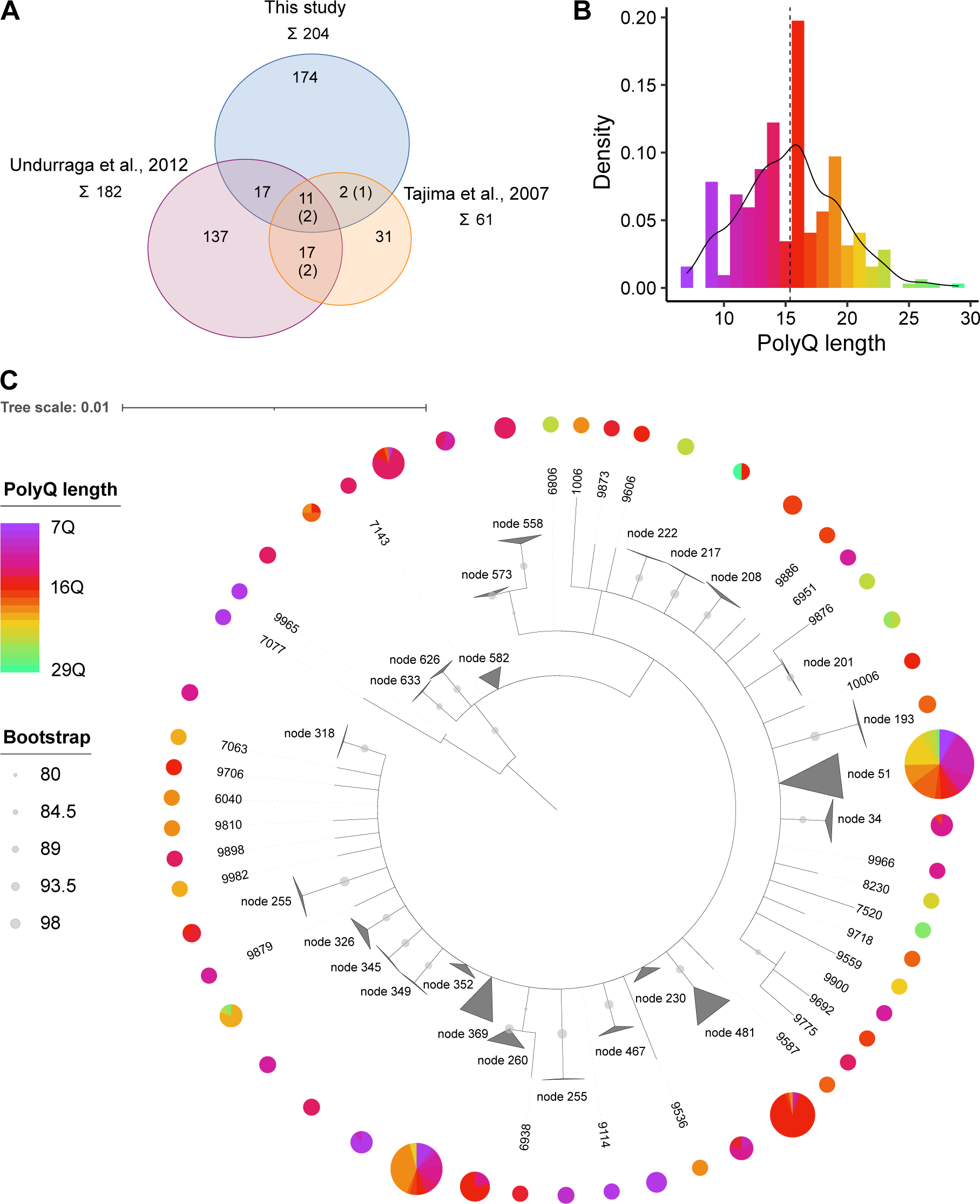
Natural variation of ELF3-polyQ in *Arabidopsis thaliana* accessions. (A) An overview of accessions with known polyQ length in ELF3. The numbers in parenthesis are the not-matching counts from different sources. In total 319 accessions included in the 1001 Genomes collection were used in this study. (B) Density plot represents the distribution of polyQ length. The dashed line represents the mean polyQ length. (C) Phylogeny of *Arabidopsis thaliana ELF3* independent of polyQ variation. Phylogenetic tree was constructed with the coding sequences (CAA repeats removed) using maximum likelihood IQ-Tree MG+F3X4 model with 10,000 replications of ultrafast bootstrap (bootstrap values >= 80 are shown as grey circles). Identical sequences are collapsed into one node. The pie chart shows the polyQ length of each leaf or node. The size of the pie chart is related to the number of leaves in each node. The accession ID, node ID, corresponding accession name, and polyQ length are listed in Supplemental Table S2.

Based on the coding sequence of *ELF3* in 319 accessions, we first tested whether *Arabidopsis thaliana* ELF3 is under any directional selection pressure. Sliding window analyses were performed for sequence polymorphism (πa/πs), as well as sequence divergence (Ka/Ks) using nine *Brassicaceae ELF3* as an interspecific group. While πa/πs refers to the genetic variation of *ELF3* within 319 accessions, Ka/Ks applies to the inter-species variation (Fay and Wu, 2003). Across the coding region of *ELF3*, few πa/πs and Ka/Ks peaks (> 1) were observed, with one Ka/Ks peak within the PrLD region, indicating that these sites may be under positive selective pressure (Supplemental Fig. S5A). The highest peak of both πa/πs and Ka/Ks was detected at the same site outside the PrLD. However, this could be explained by a relatively low synonymous substitution rate (Ks) at the site, as the overall nonsynonymous substitution rate (Ka) and nucleotide diversity (π) were very low in *ELF3* (Supplemental Fig. S5A, B). The latter suggests that apart from the polyQ variation, *ELF3* is highly conserved among *Arabidopsis thaliana* accessions. And indeed, mostly null, or negative values of Tajima’s D were detected across the coding region, with an overall value of -2.45 (*P*<0.001) (Supplemental Fig. S5C). The negative Tajima’s D indicates that *Arabidopsis thaliana ELF3* might have experienced a recent selective sweep.

Although only limited *ELF3* sequence variation was detected in *Arabidopsis thaliana* accessions, we next asked whether it might be associated with polyQ variation, which would suggest that polyQ variation could be regarded as the driver of general sequence variation within ELF3. We therefore constructed a phylogenetic tree using the obtained 319 *ELF3* sequences with the expanded CAA repeats (encoding polyQ stretch) removed. Consistent with the population genetic data (Supplemental Fig. S5), *ELF3* sequences outside of the polyQ stretch were highly conserved, as several groups of sequences were identical (shown as collapsed nodes in the phylogenetic tree in Fig. 4C). In case the length of the polyQ stretch would be a driver of ELF3 diversification within the worldwide *Arabidopsis thaliana* germplasm, other polymorphisms outside the polyQ stretch would have co-evolved and we would expect an accumulation of polyQs of similar length in specific branches of the generated phylogeny. However, we observed wide distributions of polyQ length within these collapsed nodes, suggesting that the general clustering of sequences in the phylogenetic tree was not based on the polyQ length. This indicates that even if polyQ variation might be of evolutionary relevance, it is not the driving force of *ELF3* evolution in *Arabidopsis thaliana*.

### *Arabidopsis thaliana* ELF3-polyQ variation is not likely associated with geographic origins

Based on previously published results on ELF3-polyQ function and variation (Undurraga et al., 2012; Jung et al., 2020), it has been suggested that such variation is an evolutionary adaptation to diverse latitudes and/or climates (Wilkinson and Strader, 2020; Xu et al., 2021). To test this hypothesis, we first plotted all obtained accessions according to their ELF3-polyQ length and geographic origins (coordinates) on a map focusing on European regions (where most accessions were collected). We did not detect specific distribution patterns of ELF3-polyQ length, as accessions collected from nearby sites regularly vary in polyQ length (Fig. 5A). For instance, two accessions with 26Q from Spain (ID: 9584, Supplemental Table S2) and Central Europe (ID: 7520) were both mixed with accessions with relatively short polyQ stretches in their ELF3. However, when considering all accessions, despite being weak, there is a negative correlation between polyQ length and latitude (Fig. 5B), as well as a positive correlation between polyQ length and elevation (Fig. 5C). This can be explained by the detection of accessions with long ELF3-polyQ stretches in non-European regions (Supplemental Fig. S6). For example, all four accessions from Azerbaijan had 22-23Q (ID: 9069, 9070, 9089, and 9091), one accession carrying the longest polyQ was from the Indian Ladakh plateau (29Q, ID: 8424), and one with 27Q was from Japan (ID: 7207) (Supplemental Table S2; Supplemental Fig. S6). Nevertheless, since accessions with long polyQ stretches are also present in the European region with relatively high latitude and low altitude, the overall association between polyQ variation and geographic data is not convincing.

**Fig. 5.**
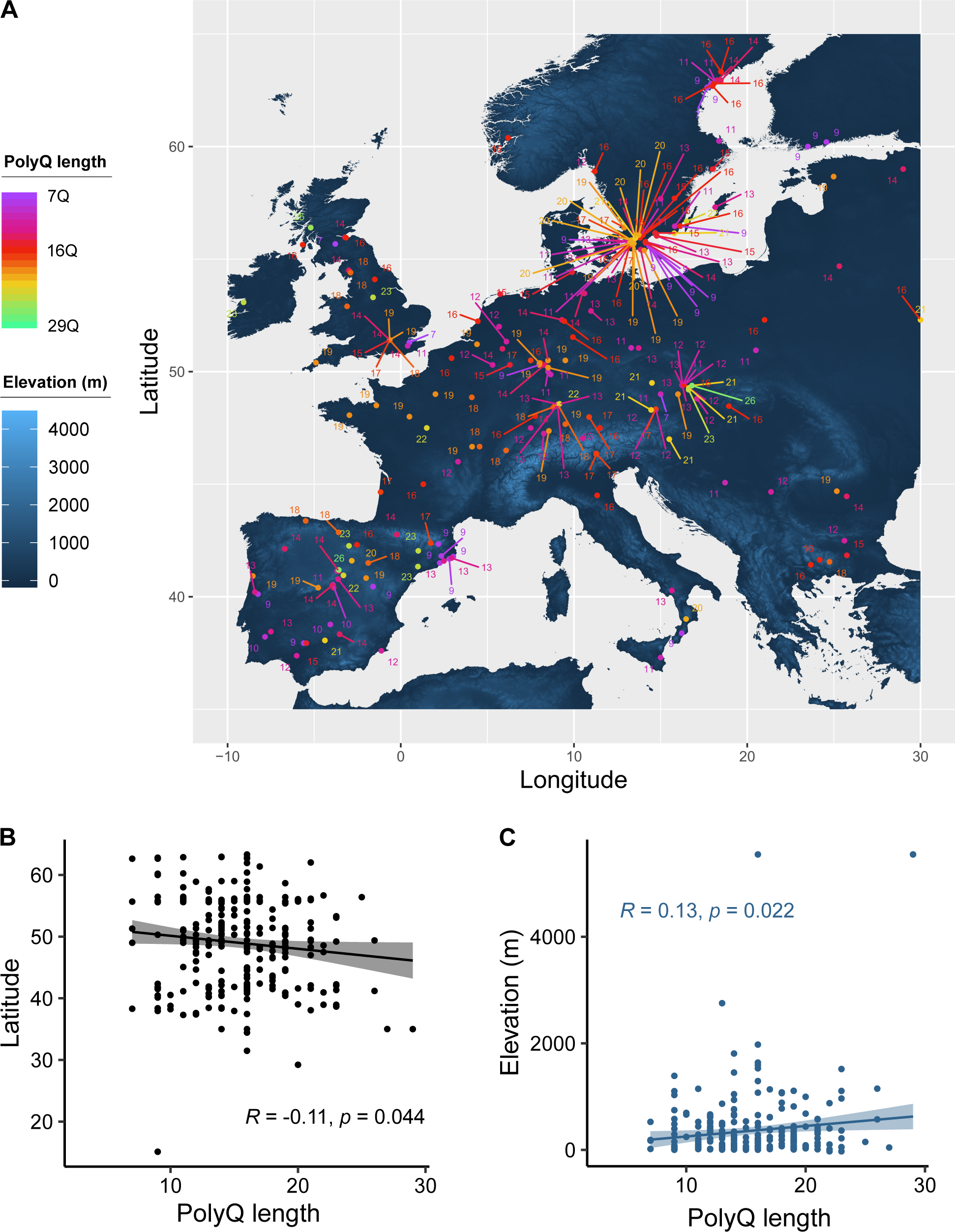
Geographic distribution of *Arabidopsis thaliana* ELF3-polyQ variation. (A) The polyQ length of each accession was mapped with the coordinates of their geographic origins. The continent is coloured based on the elevation information. The map focuses on the European region (only accessions within the area bounded by longitudes -10 to 30 and latitudes 35 to 65 are displayed), whereas the worldwide map is shown in Supplemental Fig. S6. (B, C) Pearson correlation of the polyQ length with latitude (B) or elevation (C) data. The accession ID, polyQ length, and corresponding geographic data are listed in Supplemental Table S2.

To further validate this conclusion, we investigated potential correlations between polyQ length and local climate data with a focus on temperature- and precipitation-related factors. While we detected a few weak correlations (significant, but mostly < 0.2) with selected precipitation-related parameters and a single temperature-related parameter (isothermality, ratio of diurnal variation to annual variation in temperatures, *p* = 0.014), the vast majority of parameters did not affect polyQ length (Supplemental Fig. S7). Taken together, based on the global scale of the 319 *Arabidopsis thaliana* accessions and the environmental data included in this survey, we did not observe convincing arguments for an important role of ELF3-polyQ length variation as a driver of evolutionary adaptation to local climates.

### *Arabidopsis thaliana* ELF3-polyQ variation is not associated with temperature-responsive phenotypes

As a multifunctional protein, ELF3 plays prominent roles in both circadian clock regulation and thermomorphogenesis. Previous studies reported a significant correlation of ELF3-polyQ length with circadian rhythm parameters in natural *Arabidopsis thaliana* accessions (Tajima et al., 2007) as well as transgenic lines (Undurraga et al., 2012). However, such associations were weaker regarding growth and developmental phenotypes at normal or elevated temperatures, which might depend on the genetic background of the transgenic lines (Undurraga et al., 2012; Press et al., 2016; Jung et al., 2020).

To investigate potential associations between ELF3-polyQ variation and temperature responsive phenotypes in natural *Arabidopsis thaliana* accessions, growth assays were performed under normal (20°C) and elevated (28°C shift) temperatures. Hypocotyl length was measured as a classic phenotype to represent temperature responsiveness. For the growth assays, 253 accessions were selected as a subset of the previously described 319 accessions with a similar distribution of polyQ variation (Fig. 4B; Fig. 6A). Greater and more divergent normalized hypocotyl length was observed after a temperature shift to 28°C compared to those kept at 20°C. However, the polyQ length did not correlate with normalized hypocotyl length at neither 20°C nor 28°C (Fig. 6B), nor with the temperature response of hypocotyl elongation (fold-change, Fig. 6C). Similarly, no association pattern could be detected using a three-dimensional visualization of polyQ length and normalized hypocotyl length at 20°C and 28°C (Fig. 6D). In addition, we performed correlation analysis based on previously reported flowering time data from 274 *Arabidopsis thaliana* accessions at 10°C and 16°C (Alonso-Blanco et al., 2016). However, similar to hypocotyl elongation, temperature-responsive flowering time was not associated with polyQ length (Supplemental Fig. S8). Consistent with previous reports using transgenic lines from two different genetic backgrounds (Press et al., 2016; Jung et al., 2020), these weak or absent associations suggest that potentially existing effects of polyQ length are either not prominent or masked by the genetic backgrounds, if existing at all.

**Fig. 6.**
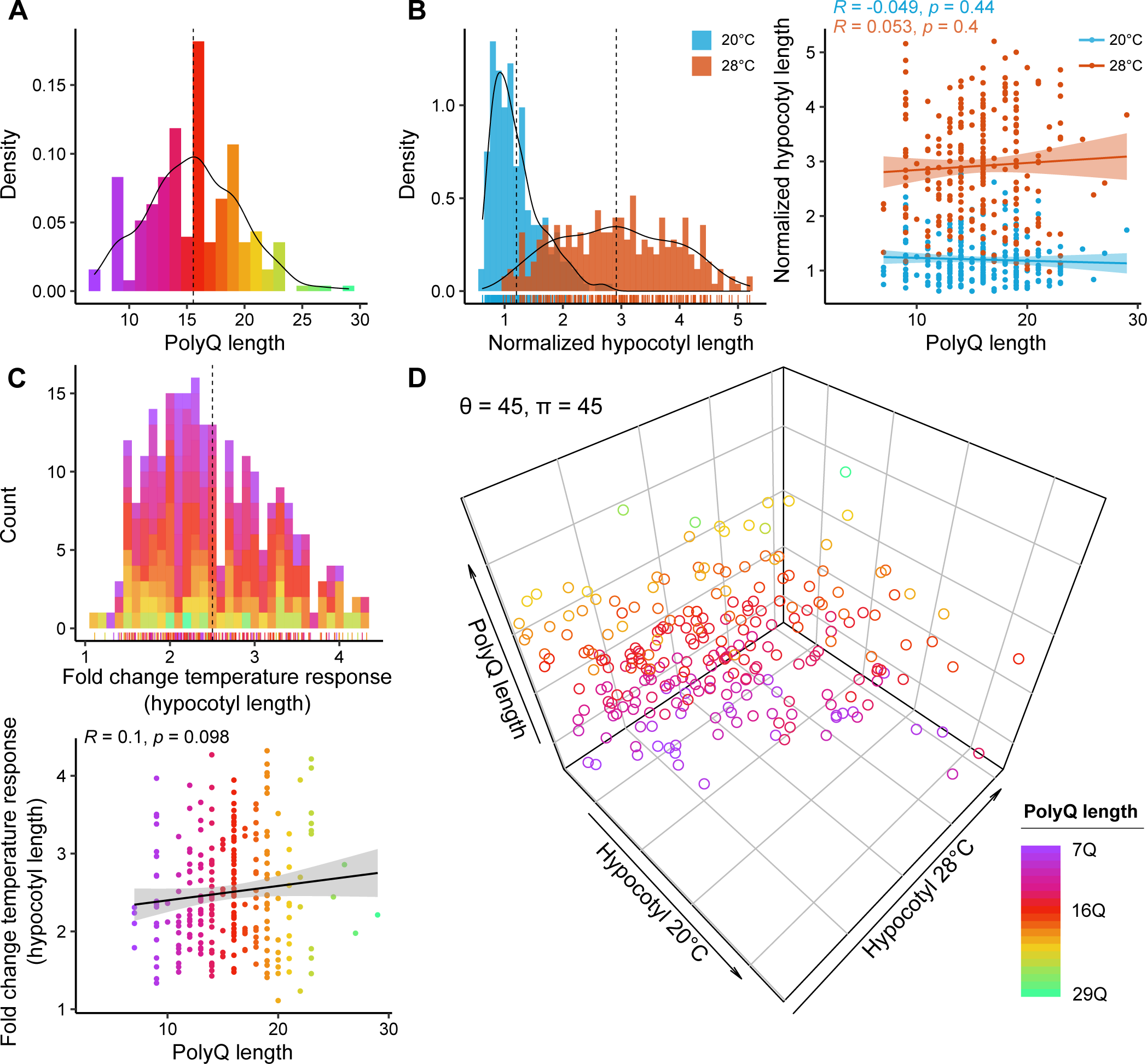
Association of ELF3-polyQ variation with temperature-responsive hypocotyl elongation. (A) Distribution of polyQ length in 253 *Arabidopsis thaliana* accessions used for growth assays (Supplemental Table S2). (B, C) Distribution of normalized hypocotyl length at 20°C or after temperature shift to 28°C (B), fold change temperature response (C), and their correlation with polyQ length. Normalized hypocotyl length represents the normalization of absolute length to median value of accession Col-0 at 20°C of each experiment. Vertical dashed lines in the distribution plots represent mean values. Arithmetic means of each accession shown as rugs below the distribution were used for Pearson correlation analysis. Colours of the stacked bars, rugs, and dots in (C) represent polyQ length as shown in (A). (D) Three-dimensional visualization of potential association among polyQ length, and normalized hypocotyl length at 20°C and 28°C. θ and π represent the rotation angles of the plot.

## Discussion

Sensing changes in ambient temperature is the first step in plant thermomorphogenesis. Among the known plant temperature sensors, the PrLD in *Arabidopsis thaliana* ELF3 mediates liquid-liquid phase separation to form aggregates at elevated temperatures (Jung et al., 2020; Hutin et al., 2023). However, it is unknown whether and how the PrLD is conserved in ELF3 orthologs across the plant kingdom. In this study, which spans genome scans of species across all major branches of the plant tree of life, we observed that the PrLD, mainly contributed by the length of polyQ, emerged and expanded most prominently in *Brassicales*. ELF3’s molecular functions in temperature-responsive aggregation are therefore not expected to be generally conserved in species outside the *Brassicales*. Although arguably limited, there is experimental evidence for lack of the ability to phase separate for the ELF3 orthologous protein in the monocot *Brachypodium distachyon* (Jung et al., 2020). Based on natural *Arabidopsis thaliana* accessions, we found that the ELF3-polyQ variation is not likely to be associated with geographic origin, climatic conditions, or classic temperature-responsive phenotypes.

Although the temperature sensing concept of ELF3-PrLD was mainly described in the model plant *Arabidopsis thaliana*, it was hypothesized that it represents an evolutionary adaptation to different climates. This hypothesis was also raised because the predicted ELF3-PrLD is either much smaller in or absent from species adapted to warmer climates such as *Solanum tuberosum* or *B. distachyon*, respectively (Jung et al., 2020). However, based on the same PrLD prediction method analyzing ELF3 homologues across the plant kingdom, we found that ELF3 from non-*Brassicales* species rarely contained a polyQ stretch or a predicted PrLD (Figs. 1-3). As replacing *Arabidopsis thaliana* ELF3 with *B. distachyon* ELF3 abolished its temperature responsiveness (Jung et al., 2020), these results suggest that the temperature sensing ability of ELF3-PrLD is only applicable to a limited number of plant species, mostly *Brassicaceae*. Nevertheless, as expected, using a prediction model based on amino acid composition like PLAAC, the probability of PrLD existence in *Brassicaceae* family significantly correlates with polyQ length which varies within the family as well as within 319 natural *Arabidopsis thaliana* accessions (Figs. 2-4; Supplemental Fig. S4). To better understand the evolutionary advantage and potential functions of polyQ variation, we closely assessed polyQ variation between *Arabidopsis thaliana* accessions. Although polyQ length presents the major sequence variation among different accessions (Supplemental Fig. S5), we failed to detect promising associations with coordinate-based geographic origins (Fig. 5). While we are aware that even nearby locations can differ drastically for selected climate factors, our data show that on the scale of this study there are no convincing arguments for an important function of ELF3-polyQ variation for an evolutionary adaptation to varying latitudes or ambient temperatures (Wilkinson and Strader, 2020; Xu et al., 2021).

Furthermore, no significant correlations between polyQ length and temperature-responsive hypocotyl or flowering phenotypes were detected in temperature assays using *Arabidopsis thaliana* accessions (Fig. 6; Supplemental Fig. S8). This could mean that some of the phenotypes (e.g., temperature-induced hypocotyl elongation) widely assessed by the thermomorphogenesis research community (and this study) are irrelevant in nature. Nevertheless, it is also consistent with a previous report which found little evidence that the polyQ stretch in transgenic lines differing in polyQ length plays a specific role in various thermal responses beyond modulating general ELF3 function (Press et al., 2016). As Col-0, the accession with the shortest polyQ stretch, has an obviously functional PrLD (Jung et al., 2020), it can be concluded that the temperature sensing properties of the ELF3-PrLD mainly depend on the ‘qualitative’ existence of polyQ, rather than its ‘quantitative’ length. However, the potential effects of polyQ length on the aggregation properties under high temperatures cannot be ruled out. For example, the detected accessions with long polyQ stretches in non-European regions may have evolutionary relevance (Supplemental Fig. S6). Such effects may be masked and/or diluted to non-detectability when global scale populations are assessed as in our study. However, when conservation of a specific polyQ length is primarily restricted to local populations and not specific climatic or geographic (e.g., latitude) factors, it is more likely that recent common ancestry of individuals of these populations is underlying this correlation and not an adaptive functional specificity conveyed by the length of polyQ.

From a physical chemistry point of view, the aggregation properties of polyQ peptides depend on both polyQ length and temperature (Walters and Murphy, 2009; Böker and Paul, 2022). The longer the peptide (= the polyQ stretch), the lower the transition temperature required for its aggregation. For example, based on computer simulations, a polyQ peptide self-aggregates at a physiological temperature (around 37°C) when its chain length is more than 25Q, whereas shorter single chains remain disordered at the same temperature (Böker and Paul, 2022). This is supported by a recent simulation study using ELF3-PrLD, concluding that increasing polyQ length promotes self-aggregation (Lindsay et al., 2023). However, whether this also applies to the thermodynamics of the entire ELF3 protein harboring polyQ and PrLD needs to be investigated at a molecular level *in planta*.

Before being revealed as a key player in temperature sensing and thermomorphogenesis, *ELF3* was initially identified as a component of the circadian clock (McWatters et al., 2000; Covington et al., 2001). Interestingly, polyQ variation in ELF3 displayed more prominent correlations with circadian rhythm parameters than with temperature-responsive phenotypes (Fig. 6; Supplemental Fig. S8) (Press et al., 2016; Jung et al., 2020). For example, polyQ length were negatively correlated with circadian phase and period in natural *Arabidopsis thaliana* accessions (Tajima et al., 2007), whereas in transgenic lines, increase (23Q) or decrease (7Q and 10Q) in polyQ length resulted in higher relative amplitude error (RAE) of circadian rhythms compared to the most frequent polyQ length (16Q) (Undurraga et al., 2012). These results suggest that the polyQ stretch (and probably the PrLD as a whole) mainly contributes to circadian clock functions, with temperature sensing being possibly only a secondary function. This hypothesis may also apply to *ELF3* itself, as the emergence and duplication of *ELF3* occurred much earlier with the other EC components (Table 1), compared to the emergence of its PrLD in *Brassicales* (Figs. 1-3).

Indeed, temperature is just one of the aspects that affect LLPS behavior (reviewed by (Xu et al., 2021). Besides environmental factors, LLPS also highly depends on the concentration and identities of macromolecules to form membraneless compartments. These compartments include cytoplasmic single-domain aggregations (e.g., purified ELF3-PrLD at high temperatures or low pH) (Jung et al., 2020; Hutin et al., 2023), as well as nuclear bodies containing photoreceptors (so-called photobodies) or circadian clock components (Ronald and Davis, 2019). These LLPS events all seem to be related to cellular localization of proteins: in a light- and temperature-dependent manner, the photoreceptor phyB reversibly accumulates in photobodies in subnuclear compartments (Yamaguchi et al., 1999; Hahm et al., 2020; Chen et al., 2022); in a time-of-day-dependent manner, circadian clock regulators such as ELF3, TOC1 (Wang et al., 2010), ELF4, and GI (Kim et al., 2007; Herrero et al., 2012) (co)localize to nuclear bodies. Interestingly, recent reports revealed that cellular localization of ELF3 responds to both ambient high temperature (Ronald et al., 2021) and light quality (Ronald et al., 2022), further suggesting that the LLPS behavior of ELF3 may not be PrLD-dependent or limited to a temperature response.

## Conclusions

Collectively, our study suggests that although presence of PrLD adds supplementary temperature sensing functions to ELF3, its regulatory role in thermomorphogenesis does not depend on this domain, and thereby its thermosensory function. Across different branches of the plant kingdom, ELF3 likely and primarily confers thermosensory-independent functions to thermomorphogenesis signaling. In that sense, the PrLD can be regarded as a lineage-specific add-on that does not significantly affect temperature responsiveness on an evolutionary scale across lineages. However, please note that these conclusions are to a large degree based on sequence predictions. Much more detailed experimental evidence than currently available is needed to either prove or disprove these hypotheses.

## Materials and methods

### Plant materials and growth conditions

Natural accessions of *Arabidopsis thaliana* obtained from Nottingham Arabidopsis Stock Centre (NASC) are listed in Supplemental Table S2. For screening of 253 accessions, seeds were surface-sterilized by washing with 70% ethanol for 3 min, and with 4% NaClO (with 0.3% TritonX) for 8 min using an orbital shaker. Seeds were then rinsed with sterile water three times for 10 min each and stratified in sterile water for 3 d at 4°C in darkness. Sterilized seeds were allowed to germinate on solid *Arabidopsis thaliana* solution (ATS) nutrient medium with 1% (w/v) sucrose (Lincoln et al., 1990). Seedlings were grown on vertically oriented plates in long days (LDs, 16 h light: 8 h dark) with 90 μmol m^−2^s^-1^ photosynthetically active radiation using white fluorescent lamps (T5 4000K). Seedlings were grown at constant 20°C for 4 d, and were either shifted to 28°C or kept at 20°C for an additional 4 d. Seedlings were imaged and the length of the hypocotyl was measured using RootDetection 0.1.2 (http://www.labutils.de/rd.html; Janitza et al., 2024). The experiments were performed separately in nine sequential batches and Col-0 (Accession ID: 6909, Supplemental Table S2) was included in each batch (*n* = 6-32). To compare the data obtained among different batches, the hypocotyl length of each accession was calculated by normalizing the absolute value to the median hypocotyl length of Col-0 at 20°C for each batch. Flowering time at 10°C (FT10) and flowering time at 16°C (FT16) data of 274 accessions were obtained from the 1001 Genomes (Alonso-Blanco et al., 2016).

### DNA sequencing

From the 1001 Genomes (https://1001genomes.org), *ELF3* coding sequences of 319 *Arabidopsis thaliana* accessions were obtained. As these sequences contained a large proportion of unknown nucleotides in *ELF3* regions encoding polyQ, polyQ variation of 115 accessions was corrected with previously published dideoxy sequencing data (Tajima et al., 2007; Undurraga et al., 2012). In addition, the PrLD regions were dideoxy sequenced and corrected in *ELF3* of the other randomly selected 204 additional accessions (Supplemental Table S2). The PrLD regions including polyQ were amplified using DreamTaq DNA polymerase (Thermo Fisher Scientific, Waltham, USA) and submitted to Eurofins Genomics (Ebersberg, Germany) for dideoxy sequencing. The PCR and sequencing primers were forward: 5’-ACAAAGGGGTGACTCGGAGA-3’ and reverse: 5’-GTCACTCCTCCCCCATCTCT-3’.

### Phylogenetic analysis

Copy numbers of ELF3, EEC, ELF4, and LUX in 42 plant species was obtained using HMMER (Finn et al., 2011) and BLASTp (Altschul et al., 1990) searches based on the *Arabidopsis thaliana* protein and coding sequences. ELF3 and EEC copies were classified using InterProScan (Jones et al., 2014). In addition, *Arabidopsis thaliana* ELF3 (AT2G25930) and EEC (AT3G21320) protein sequences were used to identify their homologous genes from available plant genomes in Phytozome v12.1, v13 (Goodstein et al., 2012), and OneKP databases (Matasci et al., 2014). In total, 434 sequences were obtained from 274 plant genomes (Supplemental Table S1). The angiosperm groups were classified based on the Angiosperm Phylogeny Website (v14, http://www.mobot.org/MOBOT/research/APweb/). Sequence alignments were performed with MUSCLE (Edgar, 2004) in AliView (Larsson, 2014) and visualized using the R package ggmsa (Zhou et al., 2022). The respective sequence alignment files are provided with the Supplemental data.

Maximum likelihood phylogenetic analysis of the sequence alignment was performed using IQ-Tree (Nguyen et al., 2015) with 10,000 replications of ultrafast bootstrap on the CIPRES Science Gateway (Miller et al., 2012). The JTT+F+R10 model was selected as the best-fit amino acid substitution model according to Bayesian Information Criterion for the phylogenetic analysis of ELF3 in green plants. The JTT+R3 model was selected for the phylogenetic analysis of *Brassicales* ELF3. All identified ELF3 and EEC sequences were subjected to localCIDER (Holehouse et al., 2017), PLAAC (Lancaster et al., 2014), and PrionW (Zambrano et al., 2015), in order to analyze intrinsically disordered protein (IDP) sequence properties and to identify probable PrLD regions. For PLAAC, the default minimum domain length of 60 amino acids was used. Background amino acids frequencies were based on *Arabidopsis thaliana* sequences. For each sequence, the COREscore (PrD score) and Log-likelihood ratio (LLR, without a hard cut-off compared to the PrD score) were retrieved to represent the probability of presence of a PrLD (Supplemental Table S1). The PrionW prediction was initially run with default settings for Q+N richness of >= 25% and a pWALTZ cut-off of 73. In order to include also the *Arabidopsis thaliana* ELF3 sequence, the PrionW restrictions were relaxed with Q+N richness set to 18% and pWALTZ cut-off set to 68. To generate a phylogenetic tree of *Arabidopsis thaliana ELF3* independent of polyQ, the CCA repeats (polyQ stretch) and the stop codon (as well as the sequence after a premature stop codon in one accession, ID: 9089, Supplemental Table S2) were removed from the corrected 319 *ELF3* coding sequences. Sequence alignment and phylogenetic analysis were performed as described above. The MG+F3X4 model was selected as the best-fit codon model. Phylogenetic trees were visualized and annotated in iTOL (Letunic and Bork, 2007).

### Population genetic analysis

Sequence polymorphism (πa/πs), nucleotide diversity (π), and Tajima’s D (Tajima, 1989) of *ELF3* were calculated among 319 *Arabidopsis thaliana* accessions, as well as sequence divergence (Ka/Ks) of *ELF3* between *Arabidopsis thaliana* and other *Brassicaceaes*, using sliding window analyses (width: 30 bp, step: 3 bp) in DnaSP v6 (Rozas et al., 2017). The *ELF3* sequences of nine *Brassicaceae* species (*Arabidopsis lyrata*, *Arabidopsis halleri*, *Brassica oleracea*, *Boechera stricta*, *Capsella rubella*, *Crambe hispanica*, *Descurainia sophioides*, *Eutrema salsugineum*, *and Thlaspi arvense*) were used as an interspecific group for Ka/Ks analysis.

### Association analysis

Geographic distribution of *Arabidopsis thaliana* accessions was mapped based on the coordinates using R packages geodata and ggrepel. The local environment data of 317 accessions were obtained from the Arabidopsis CLIMtools (Ferrero-Serrano and Assmann, 2019). Pairwise correlation analysis was performed with the polyQ length and visualized using the R package corrplot. Distributions and Pearson correlations of polyQ length and phenotypic data were computed and visualized using packages ggpubr and plot3D in R.

## Supporting information

Alignments

## Acknowledgements

The authors thank Wolfgang Paul (MLU Halle) for explanation of the biophysical features of polyQ stretches in peptides. Funding of this work was provided by grants from the European Social Fund and the Federal State of Saxony-Anhalt (International Graduate School AGRIPOLY - Determinants of Plant Performance, grant no. ZS/2016/08/80644) to MQ.

## Supplemental data

**Supplemental Table S1.** PrLD prediction of identified ELF3 and EEC homologues in 274 plant genomes.

**Supplemental Table S2.** PolyQ length and temperature responsive phenotypes of *Arabidopsis thaliana* accessions used in this study.

**Supplemental Fig. S1.**
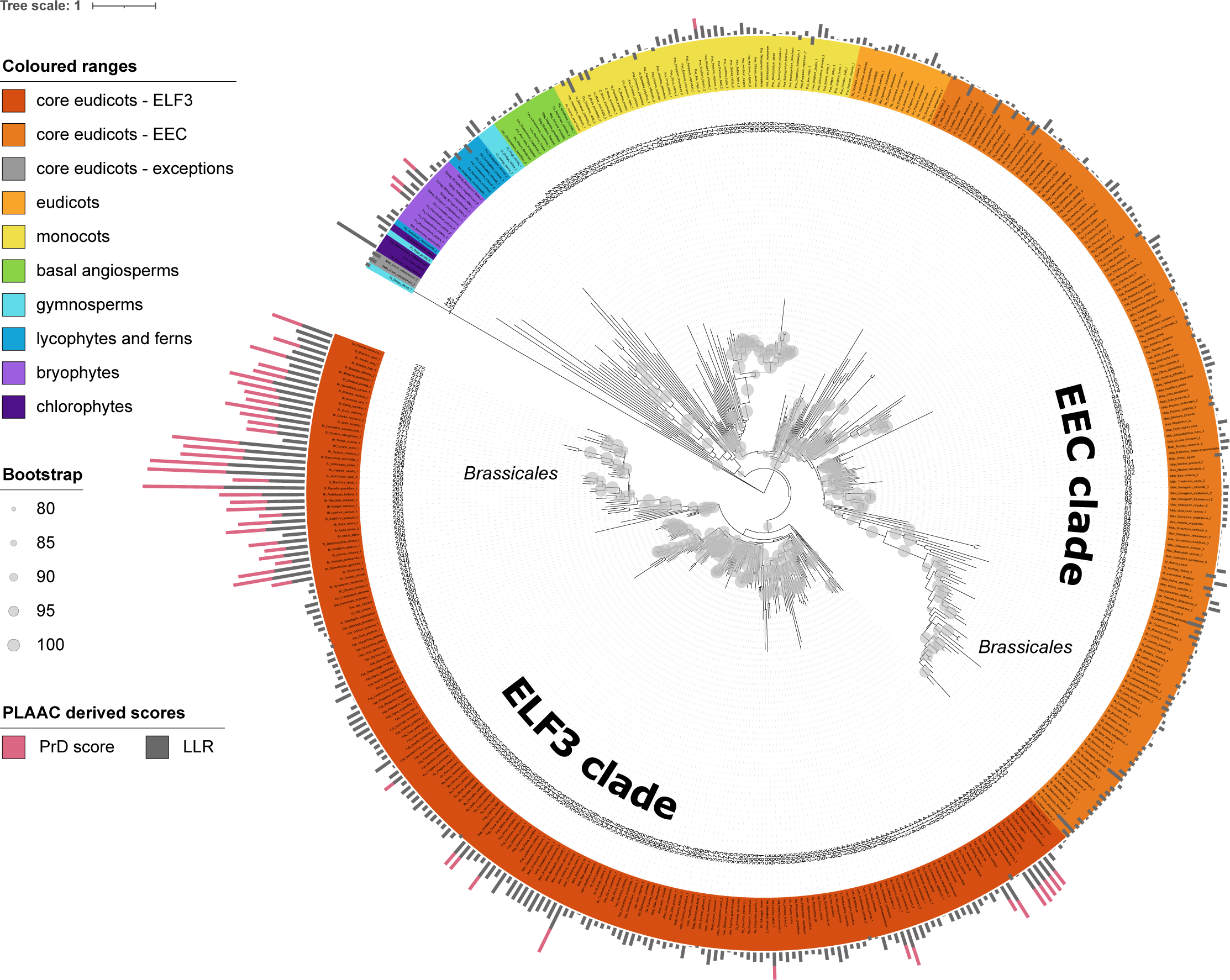
Phylogeny of ELF3 and EEC across the plant kingdom (full tree). Phylogenetic tree was constructed with the full-length amino acid sequences obtained from 274 plant genomes, using maximum likelihood IQ-Tree JTT+F+R10 model with 10,000 replications of ultrafast bootstrap (bootstrap values >= 80 are shown as grey circles). ELF3 and EEC clades of core eudicots are marked based on the position of *Arabidopsis thaliana* ELF3 and EEC, respectively. The labels are coloured according to species group and clade. PLAAC derived scores are shown as stacked bar charts outside of the tree. Leaf names and scores are listed corresponding to the branch ID in Supplemental Table S1.

**Supplemental Fig. S2.**
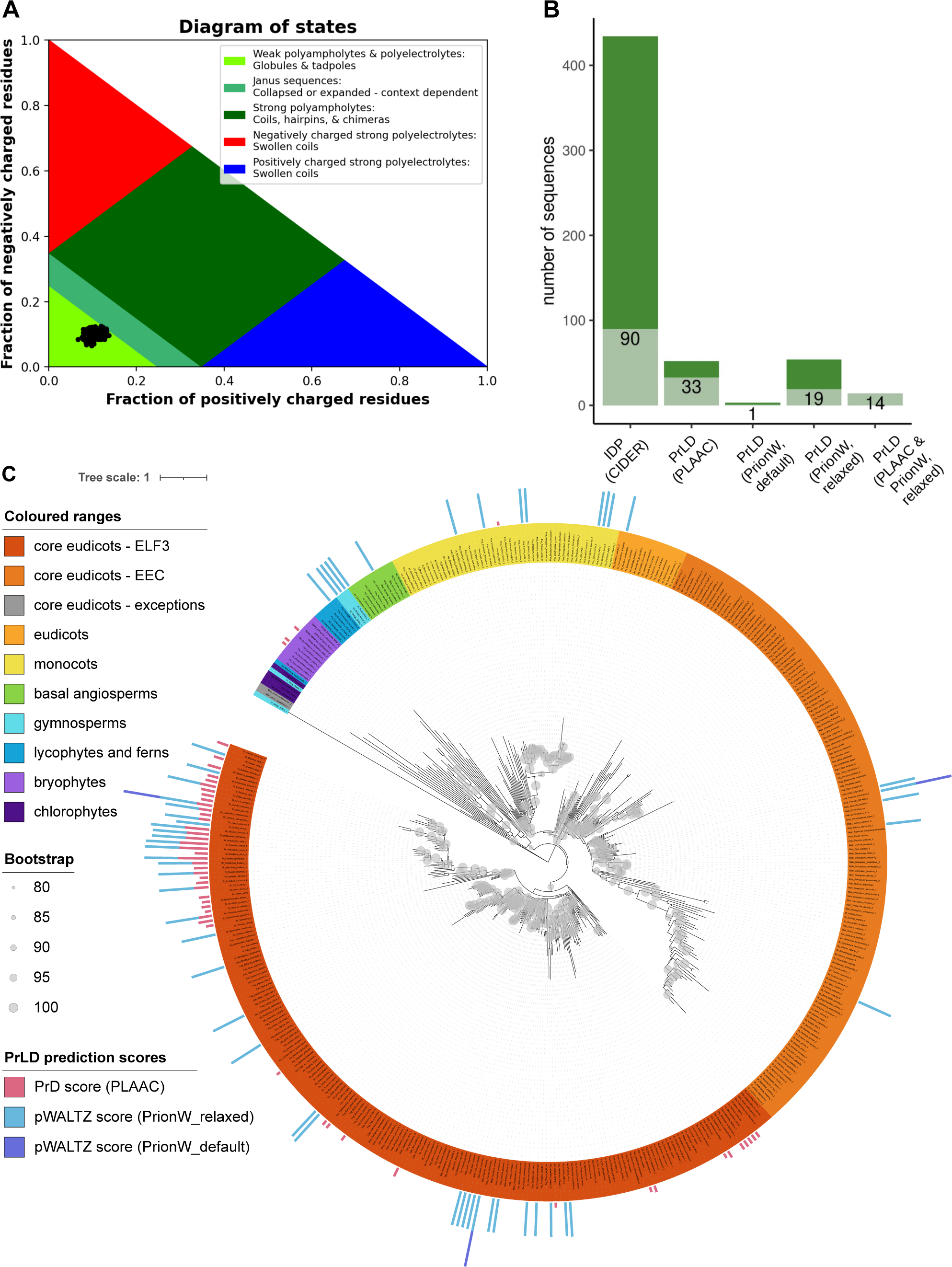
Complementary sequence analyses of ELF3/EEC homologues. (A) CIDER diagram of states for all 434 ELF3/EEC homologues. Based on their fractions of positively and negatively charged residues, the analyzed sequences (black circles) are partitioned into one of five different conformational classes (Holehouse et al., 2017). (B) Comparison of PrLD prediction results by PLAAC, PrionW_default, and PrionW_relaxed. The grey bars indicate the fraction of *Brassicales* species. (C) Comparison of PrLD prediction scores by PLAAC, Prion_default, and PrionW_relaxed in the context of ELF3/EEC phylogeny. The phylogenetic tree corresponds to Supplemental Fig. S1. PLAAC and PrionW derived scores are shown as stacked bar charts outside of the tree. Leaf names and scores are listed corresponding to the branch ID in Supplemental Table S1.

**Supplemental Fig. S3.**
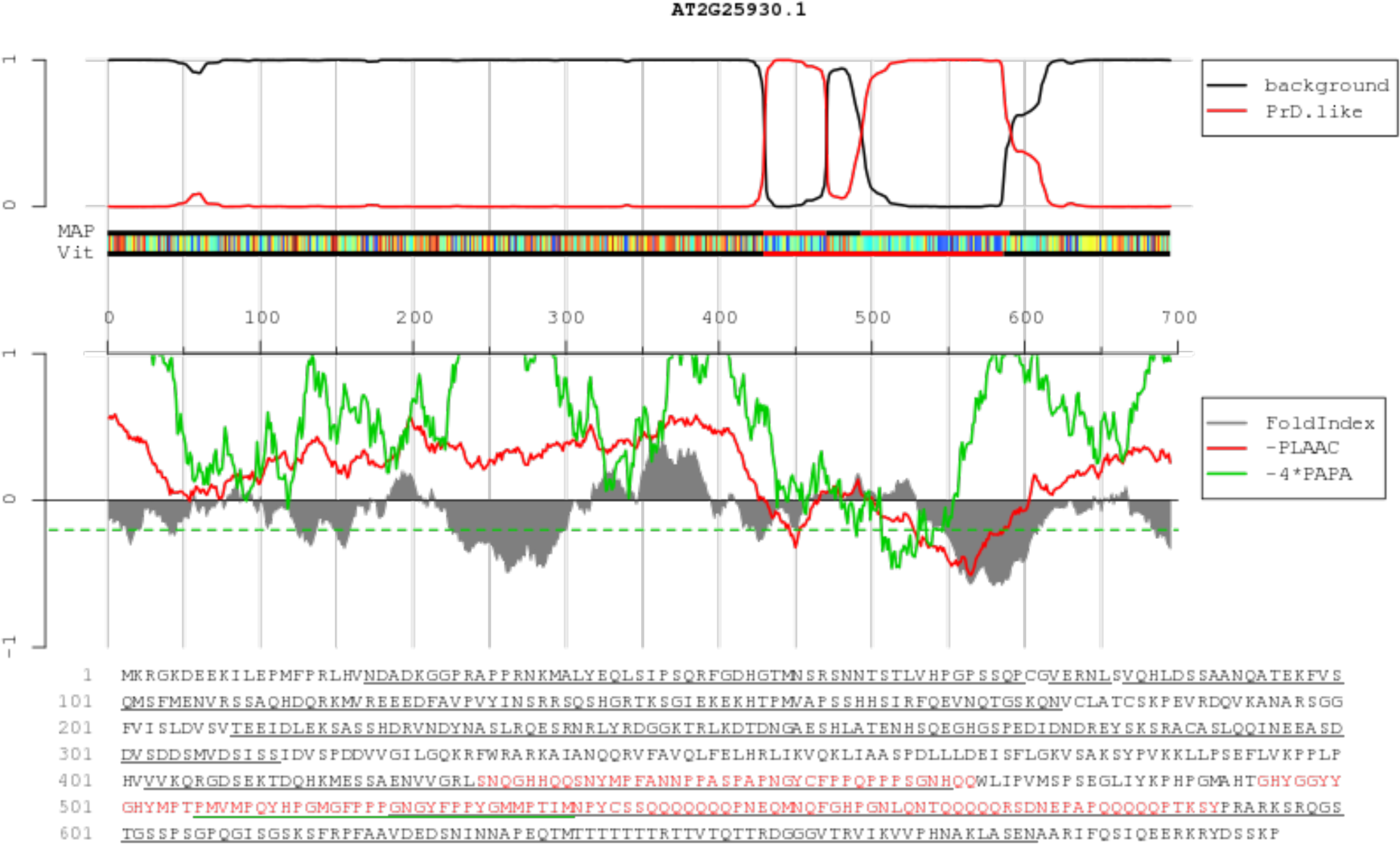
The PrLD of *Arabidopsis thaliana* ELF3. The visual output of the PLAAC analysis (Alberti et al., 2009) of *Arabidopsis thaliana* ELF3 with a default minimum domain length of 60 amino acids consists of three corresponding plots and the annotated amino acid sequence. On top, the sliding averages of per-residue log-likelihood ratios for the prion-like (red line) and background state (black line) are plotted. The next panel shows the probability of each residue belonging to the HMM state ‘PrD.like’ (red) and ‘background’ (black); the tracks ‘MAP’ and ‘Vit’ illustrate the Maximum a Posteriori and the Viterbi parses of the ELF3 protein into these two states. The lower panel shows sliding averages over a window of width 60 of predicted disorder (grey) as FoldIndex (Prilusky et al., 2005). The -PLAAC track (red) are these sliding averages scaled by using base -4 and reserved in sign. The green track is the re-implementation of PAPA (Toombs et al., 2010; Toombs et al., 2012) which is multiplied by -4 so that lower scores are more predictive of prion propensity, and so that the range is more comparable to the other tracks. A dashed green line represents a similarity rescaled version of the cutoff PAPA > 0.05 (Lancaster et al., 2014).

**Supplemental Fig. S4.**
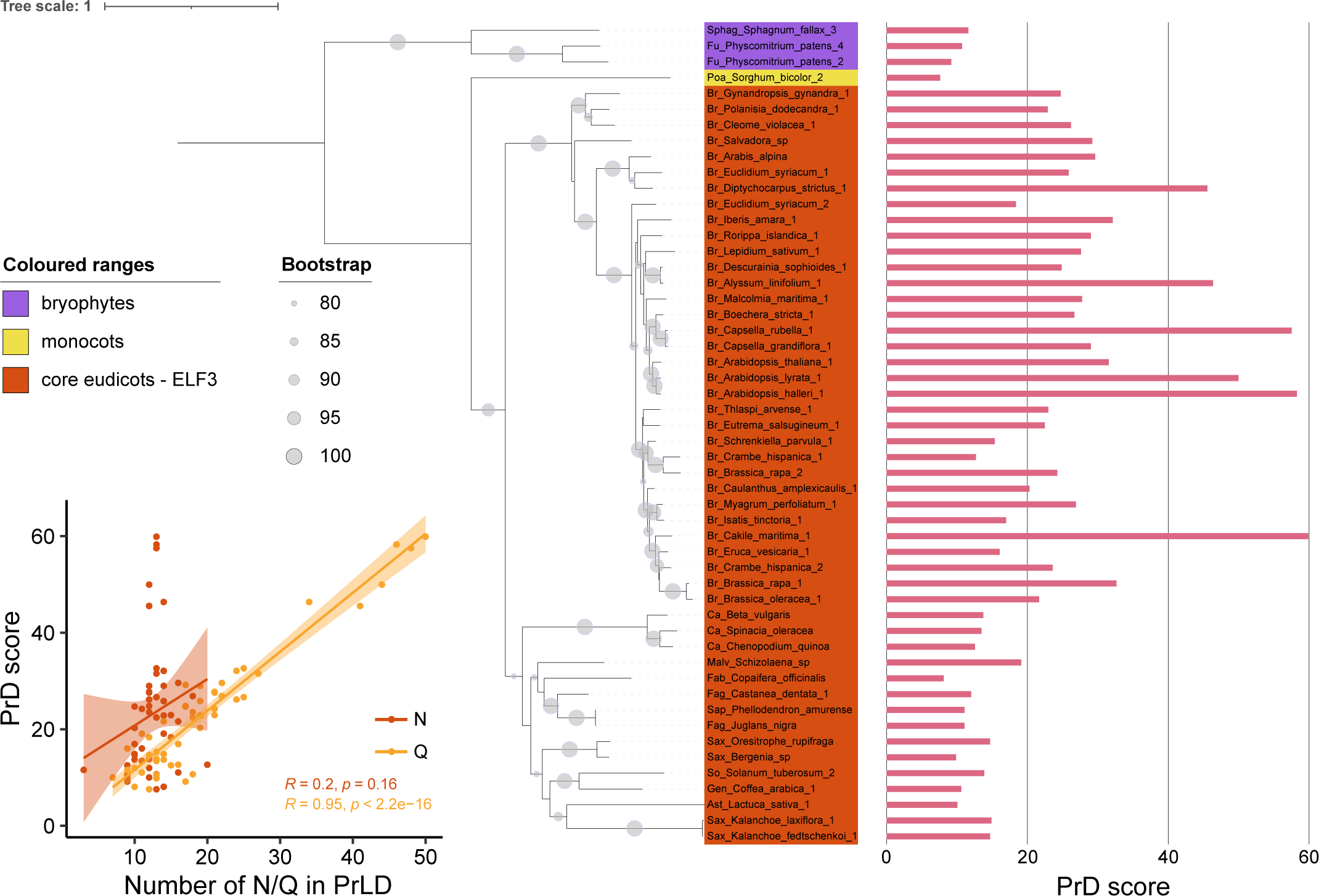
ELF3-PrLD is mainly contributed by a polyQ stretch. The phylogenetic tree was constructed including all full-length amino acid sequences with positive PrD scores, using maximum likelihood IQ-Tree JTT+R3 model with 10,000 replications of ultrafast bootstrap (bootstrap values >= 80 are shown as grey circles). PrD score is shown as bar chart. Pearson correlation of PrD score and the number of asparagine (N) or glutamine (Q) in the region.

**Supplemental Fig. S5.**
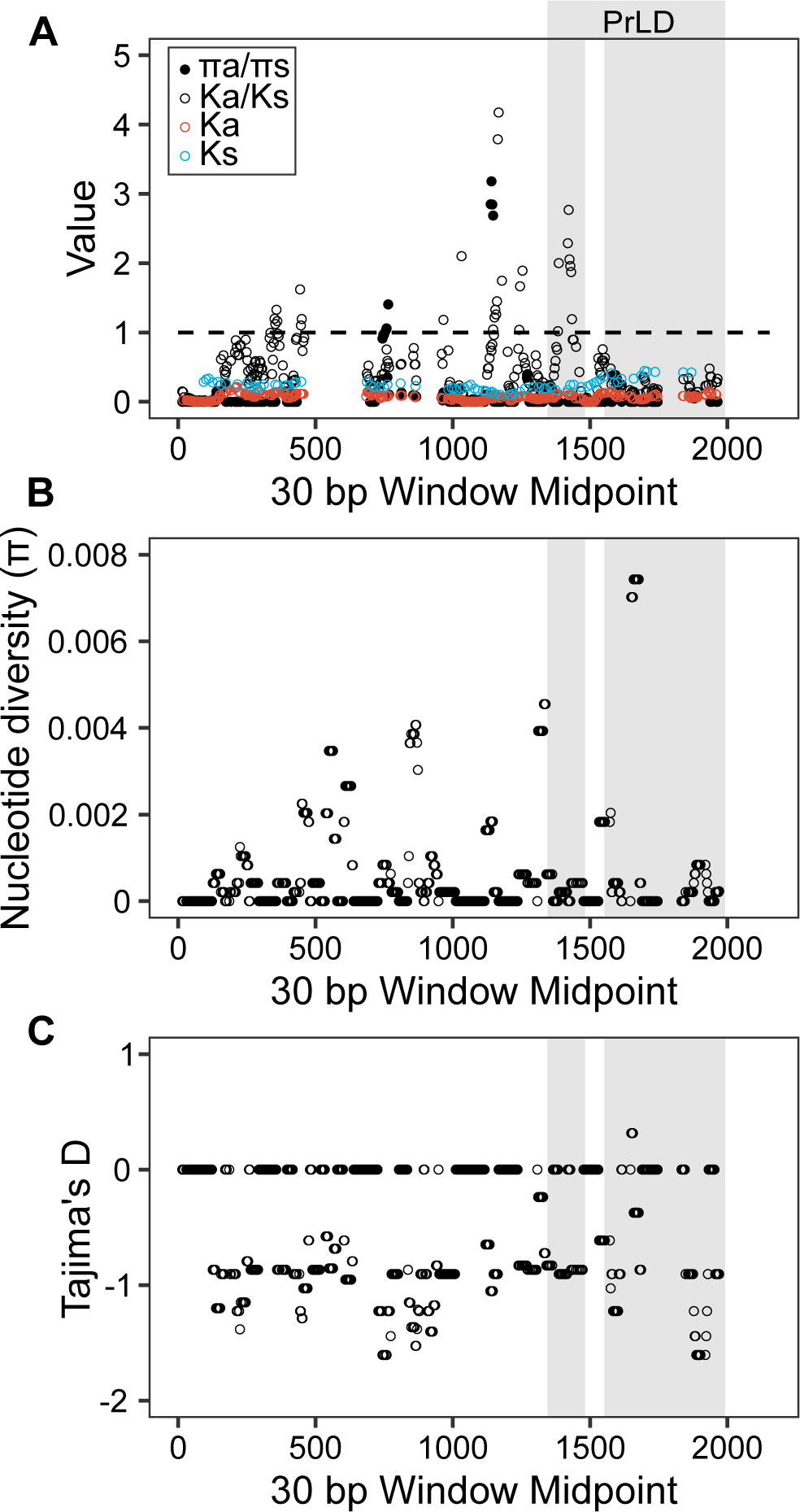
Population genetic signatures of *Arabidopsis thaliana ELF3*. (A-C) Sequence polymorphism and divergence (A), nucleotide diversity (B), and Tajima’s D (C) of full-length ELF3 coding sequence were calculated from 319 *Arabidopsis thaliana* accessions (Supplemental Table S2) using sliding window analyses (width: 30 bp, step: 3 bp). The ELF3 sequences of nine *Brassicaceae* species (*Arabidopsis lyrata*, *Arabidopsis halleri*, *Brassica oleracea*, *Boechera stricta*, *Capsella rubella*, *Crambe hispanica*, *Descurainia sophioides*, *Eutrema salsugineum*, *and Thlaspi arvense*) were used as an interspecific group for Ka/Ks analysis. Shaded areas represent the predicted regions encoding PrLD (Supplemental Fig. S3), based on the sequence alignment using *Arabidopsis thaliana* ELF3.

**Supplemental Fig. S6.**
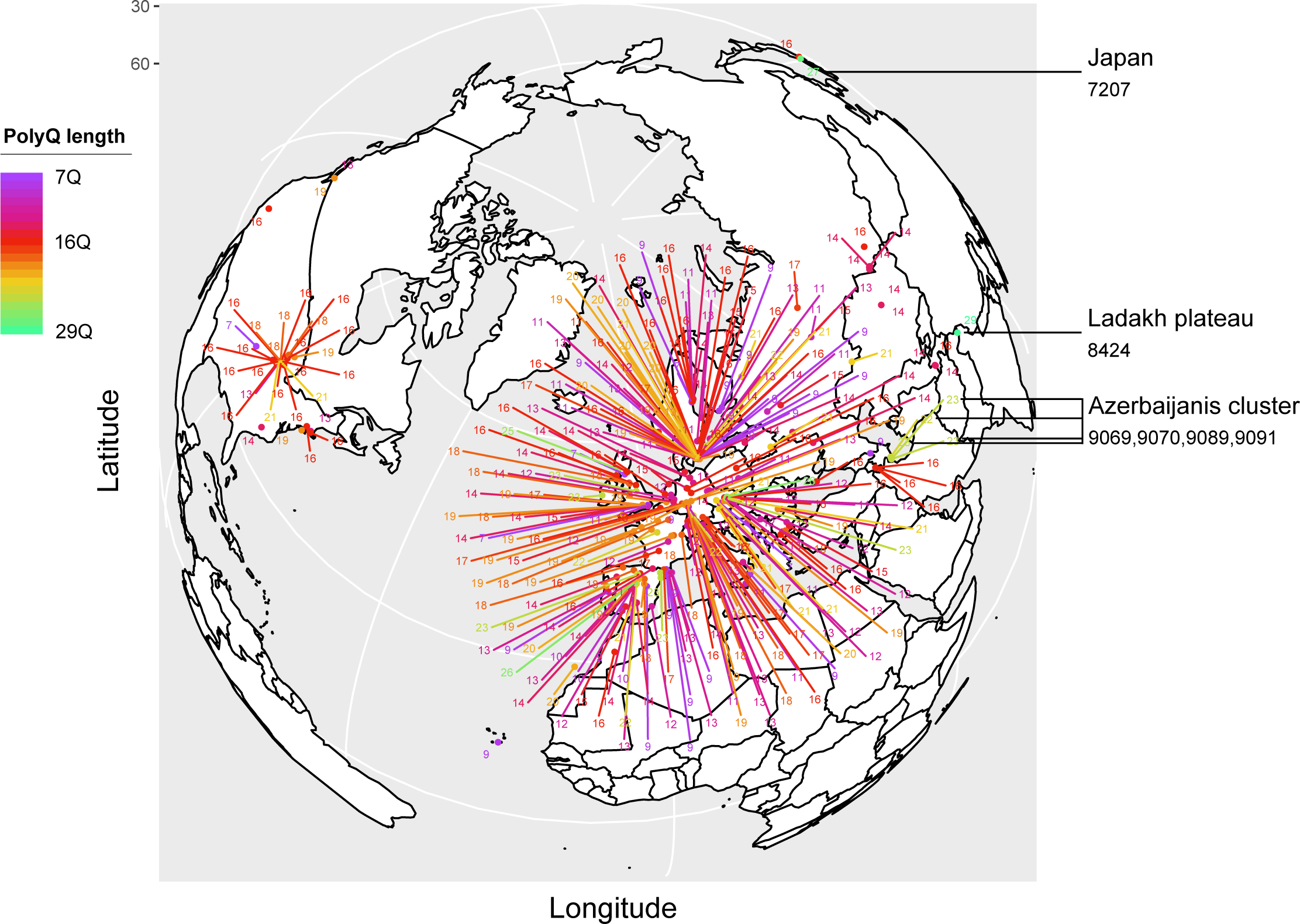
Worldwide distribution of *Arabidopsis thaliana* ELF3-polyQ variation. 319 *Arabidopsis thaliana* accessions (Supplemental Table S2) were plotted on a world map with corresponding polyQ length. Accessions with special focus are marked with accession ID and their geographic origins.

**Supplemental Fig. S7.**
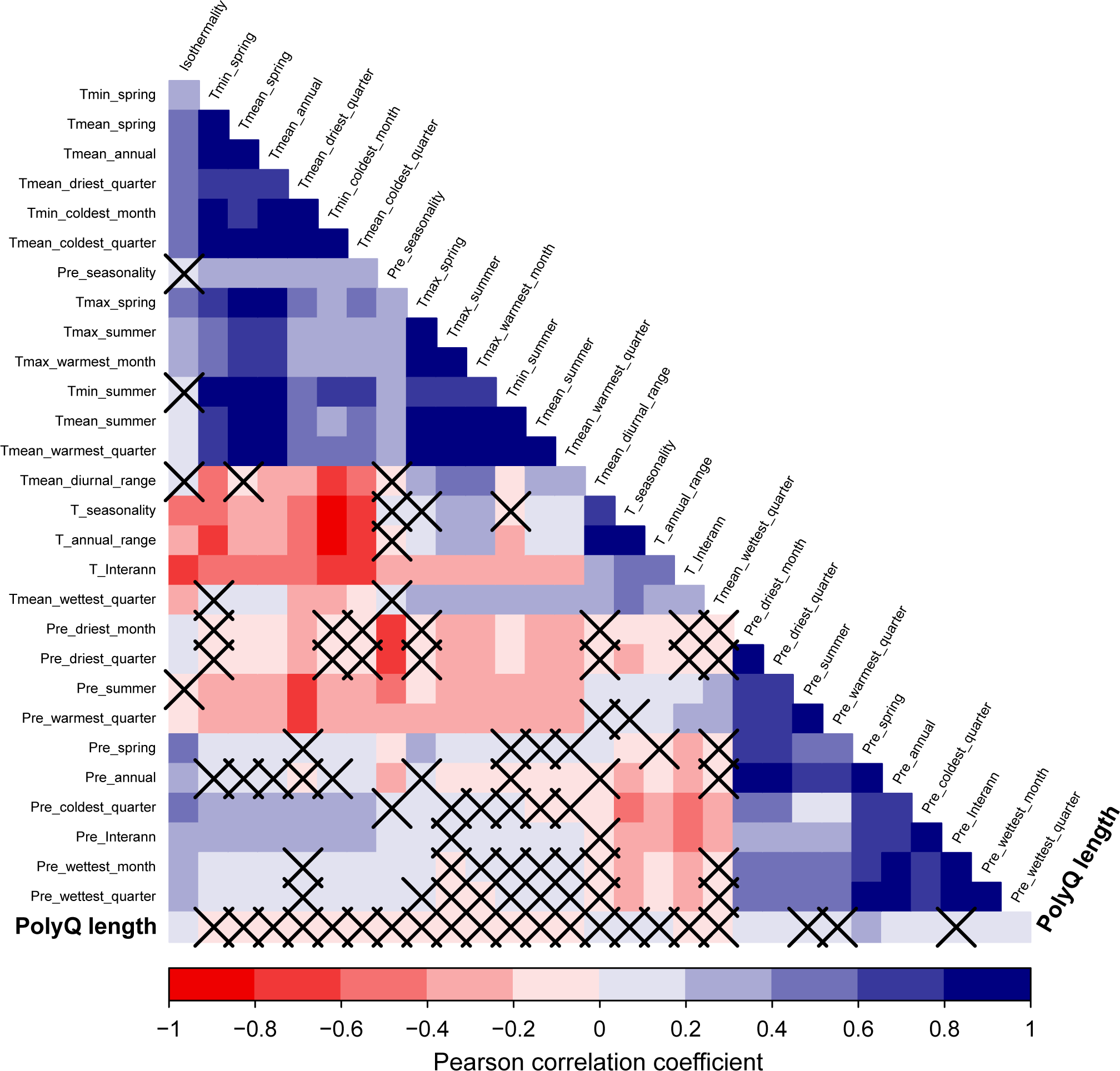
*Arabidopsis thaliana* ELF3-polyQ length is not associated with local environmental data. Pairwise correlation was determined between polyQ length, and temperature (T) or precipitation (Pre) related parameters. Pearson correlation coefficients were tested for significance and only significant coefficients with *P* < 0.05 are not crossed. The obtained CHELSA (Climatologies at high resolution for the earth’s land surface areas) climate data is described at gramene.org/CLIMtools/arabidopsis_v2.0/environments.html.

**Supplemental Fig. S8.**
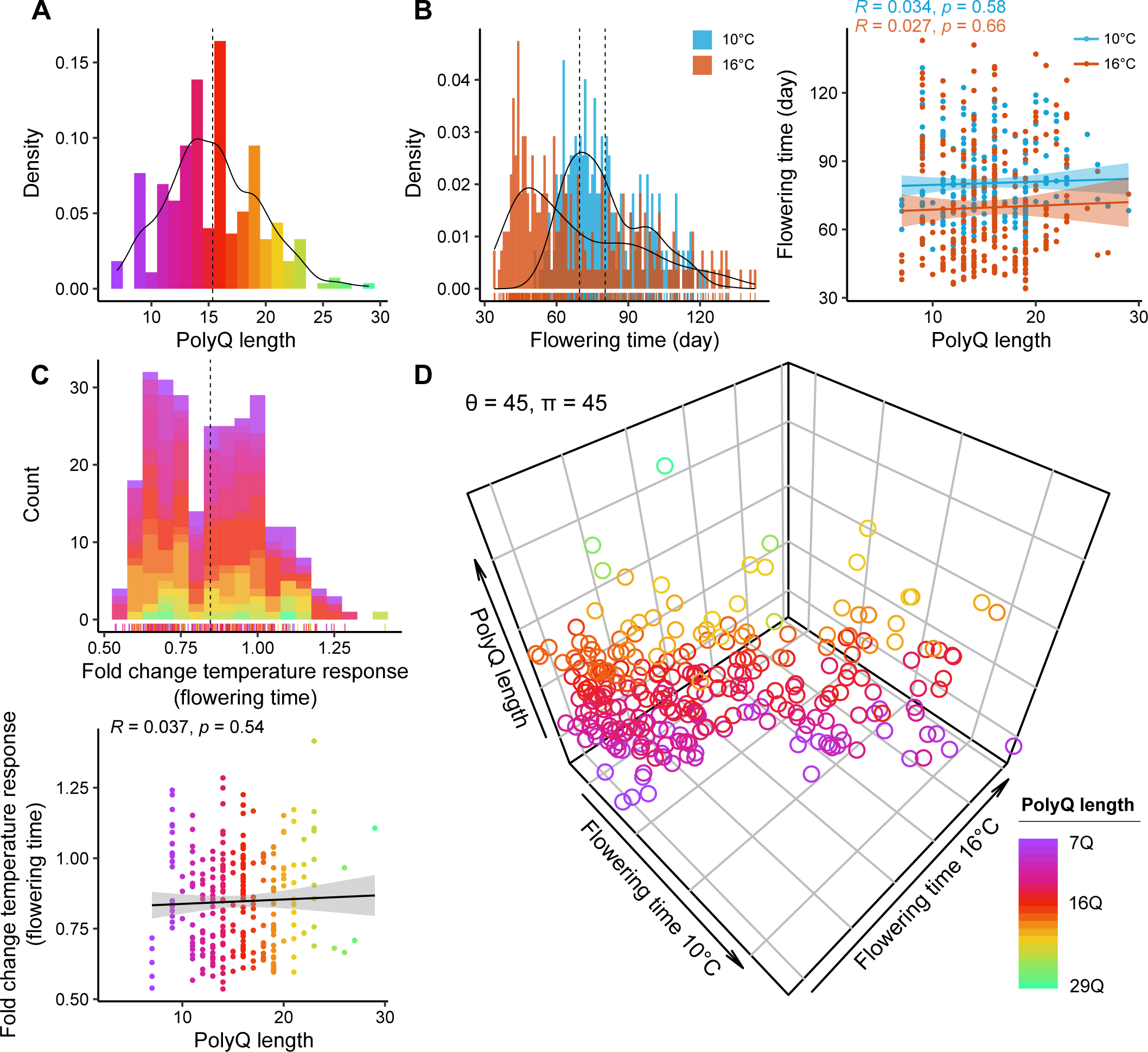
Association of ELF3-polyQ variation with temperature-responsive flowering. (A) Distribution of polyQ length in 274 *Arabidopsis thaliana* accessions used for the analysis (Supplemental Table S2). (B, C) Distribution of flowering time at 10°C or 16°C (B), fold change temperature response (C), and their correlation with polyQ length. Vertical dashed lines in the distribution plots represent mean values. Colours of the stacked bars, rugs, and dots in (C) represent polyQ length as shown in (A). (D) Three-dimensional visualization of potential association among polyQ length, and flowering time at 10°C and 16°C. θ and π represent the rotation angles of the plot.

**Alignment file 1.** Amino acid sequence alignment of 434 ELF3/EEC homologues as basis for the phylogenetic tree in Fig. 1, S1, S2C (FASTA formatted)

**Alignment file 2.** Amino acid sequence alignment of 69 selected ELF3/EEC homologues in Fig. 2 (FASTA formatted)

**Alignment file 3.** Amino acid sequence alignment of 52 selected ELF3/EEC homologues as basis for for the phylogenetic tree in Fig. 3 (FASTA formatted)

**Alignment file 4.** ELF3 coding sequence alignment of 319 *Arabidopsis thaliana* accessions as basis for the phylogenetic tree in Fig. 4C (FASTA formatted)

## Notes

### Competing Interest Statement

The authors have declared no competing interest.

### Summary of Updates

We have added the results of additional sequence motif prediction software to substantiate the previously reported data.

## References

Adams S, Manfield I, Stockley P, Carré IA (2015) Revised morning loops of the Arabidopsis circadian clock based on analyses of direct regulatory interactions. PLoS One 10: e0143943

Alberti S, Halfmann R, King O, Kapila A, Lindquist S (2009) A systematic survey identifies prions and illuminates sequence features of prionogenic proteins. Cell 137: 146–158

Alonso-Blanco C, Andrade J, Becker C, Bemm F, Bergelson J, Borgwardt KM, Cao J, Chae E, Dezwaan TM, Ding W (2016) 1,135 genomes reveal the global pattern of polymorphism in Arabidopsis thaliana. Cell 166: 481–491

Altschul SF, Gish W, Miller W, Myers EW, Lipman DJ (1990) Basic local alignment search tool. Journal of molecular biology 215: 403–410

Alvarez M, Tranquilli G, Lewis S, Kippes N, Dubcovsky J (2016) Genetic and physical mapping of the earliness per se locus Eps-A m 1 in Triticum monococcum identifies EARLY FLOWERING 3 (ELF3) as a candidate gene. Functional & Integrative Genomics 16: 365–382

Alvarez MA, Li C, Lin H, Joe A, Padilla M, Woods DP, Dubcovsky J (2023) EARLY FLOWERING 3 interactions with PHYTOCHROME B and PHOTOPERIOD1 are critical for the photoperiodic regulation of wheat heading time. PLoS Genetics 19: e1010655

Andrade L, Lu Y, Cordeiro A, Costa JM, Wigge PA, Saibo NJ, Jaeger KE (2022) The evening complex integrates photoperiod signals to control flowering in rice. Proceedings of the National Academy of Sciences 119: e2122582119

Anwer MU, Davis A, Davis SJ, Quint M (2020) Photoperiod sensing of the circadian clock is controlled by EARLY FLOWERING 3 and GIGANTEA. The Plant Journal 101: 1397–1410

Blackman BK (2017) Changing responses to changing seasons: natural variation in the plasticity of flowering time. Plant physiology 173: 16–26

Böker A, Paul W (2022) Thermodynamics and Conformations of Single Polyalanine, Polyserine, and Polyglutamine Chains within the PRIME20 Model. The Journal of Physical Chemistry B 126: 7286–7297

Box MS, Huang BE, Domijan M, Jaeger KE, Khattak AK, Yoo SJ, Sedivy EL, Jones DM, Hearn TJ, Webb AA (2015) ELF3 controls thermoresponsive growth in Arabidopsis. Current biology 25: 194–199

Bu T, Lu S, Wang K, Dong L, Li S, Xie Q, Xu X, Cheng Q, Chen L, Fang C (2021) A critical role of the soybean evening complex in the control of photoperiod sensitivity and adaptation. Proceedings of the National Academy of Sciences 118: e2010241118

Challinor AJ, Watson J, Lobell DB, Howden S, Smith D, Chhetri N (2014) A meta-analysis of crop yield under climate change and adaptation. Nature climate change 4: 287–291

Chen D, Lyu M, Kou X, Li J, Yang Z, Gao L, Li Y, Fan L-m, Shi H, Zhong S (2022) Integration of light and temperature sensing by liquid-liquid phase separation of phytochrome B. Molecular Cell 82: 3015–3029. e3016

Covington MF, Panda S, Liu XL, Strayer CA, Wagner DR, Kay SA (2001) ELF3 modulates resetting of the circadian clock in Arabidopsis. The Plant Cell 13: 1305–1316

Crawford AJ, McLachlan DH, Hetherington AM, Franklin KA (2012) High temperature exposure increases plant cooling capacity. Current Biology 22: R396–R397

Delker C, Sonntag L, James GV, Janitza P, Ibanez C, Ziermann H, Peterson T, Denk K, Mull S, Ziegler J (2014) The DET1-COP1-HY5 pathway constitutes a multipurpose signaling module regulating plant photomorphogenesis and thermomorphogenesis. Cell reports 9: 1983–1989

Delker C, van Zanten M, Quint M (2017) Thermosensing enlightened. Trends in Plant Science 22: 185–187

Edgar RC (2004) MUSCLE: multiple sequence alignment with high accuracy and high throughput. Nucleic acids research 32: 1792–1797

Ejaz M, von Korff M (2017) The genetic control of reproductive development under high ambient temperature. Plant Physiology 173: 294–306

Ezer D, Jung J-H, Lan H, Biswas S, Gregoire L, Box MS, Charoensawan V, Cortijo S, Lai X, Stöckle D (2017) The evening complex coordinates environmental and endogenous signals in Arabidopsis. Nature plants 3: 1–12

Fan H-C, Ho L-I, Chi C-S, Chen S-J, Peng G-S, Chan T-M, Lin S-Z, Harn H-J (2014) Polyglutamine (PolyQ) diseases: genetics to treatments. Cell transplantation 23: 441–458

Fang X, Han Y, Liu M, Jiang J, Li X, Lian Q, Xie X, Huang Y, Ma Q, Nian H (2021) Modulation of evening complex activity enables north-to-south adaptation of soybean. Science China Life Sciences 64: 179–195

Farré EM, Harmer SL, Harmon FG, Yanovsky MJ, Kay SA (2005) Overlapping and distinct roles of PRR7 and PRR9 in the Arabidopsis circadian clock. Current Biology 15: 47–54

Faure S, Turner AS, Gruszka D, Christodoulou V, Davis SJ, von Korff M, Laurie DA (2012) Mutation at the circadian clock gene EARLY MATURITY 8 adapts domesticated barley (Hordeum vulgare) to short growing seasons. Proceedings of the National Academy of Sciences 109: 8328–8333

Fay JC, Wu C-I (2003) Sequence divergence, functional constraint, and selection in protein evolution. Annual review of genomics and human genetics 4: 213–235

Ferrero-Serrano Á, Assmann SM (2019) Phenotypic and genome-wide association with the local environment of Arabidopsis. Nature Ecology & Evolution 3: 274–285

Finn RD, Clements J, Eddy SR (2011) HMMER web server: interactive sequence similarity searching. Nucleic acids research 39: W29–W37

Ford B, Deng W, Clausen J, Oliver S, Boden S, Hemming M, Trevaskis B (2016) Barley (Hordeum vulgare) circadian clock genes can respond rapidly to temperature in an EARLY FLOWERING 3-dependent manner. Journal of Experimental Botany 67: 5517–5528

Franklin KA, Lee SH, Patel D, Kumar SV, Spartz AK, Gu C, Ye S, Yu P, Breen G, Cohen JD (2011) Phytochrome-interacting factor 4 (PIF4) regulates auxin biosynthesis at high temperature. Proceedings of the National Academy of Sciences 108: 20231–20235

Gil-Garcia M, Iglesias V, Pallarès I, Ventura S (2021) Prion-like proteins: from computational approaches to proteome-wide analysis. FEBS open bio 11: 2400–2417

Goodstein DM, Shu S, Howson R, Neupane R, Hayes RD, Fazo J, Mitros T, Dirks W, Hellsten U, Putnam N (2012) Phytozome: a comparative platform for green plant genomics. Nucleic acids research 40: D1178–D1186

Hahm J, Kim K, Qiu Y, Chen M (2020) Increasing ambient temperature progressively disassembles Arabidopsis phytochrome B from individual photobodies with distinct thermostabilities. Nature communications 11: 1660

Harrison PM, Gerstein M (2003) A method to assess compositional bias in biological sequences and its application to prion-like glutamine/asparagine-rich domains in eukaryotic proteomes. Genome biology 4: 1–14

Herrero E, Kolmos E, Bujdoso N, Yuan Y, Wang M, Berns MC, Uhlworm H, Coupland G, Saini R, Jaskolski M (2012) EARLY FLOWERING4 recruitment of EARLY FLOWERING3 in the nucleus sustains the Arabidopsis circadian clock. The Plant Cell 24: 428–443

Holehouse AS, Das RK, Ahad JN, Richardson MO, Pappu RV (2017) CIDER: resources to analyze sequence-ensemble relationships of intrinsically disordered proteins. Biophysical journal 112: 16–21

Hsu PY, Devisetty UK, Harmer SL (2013) Accurate timekeeping is controlled by a cycling activator in Arabidopsis. Elife 2: e00473

Huang H, Nusinow DA (2016) Into the evening: Complex interactions in the Arabidopsis circadian clock. Trends in Genetics 32: 674–686

Hutin S, Kumita JR, Strotmann VI, Dolata A, Ling WL, Louafi N, Popov A, Milhiet P-E, Blackledge M, Nanao MH (2023) Phase separation and molecular ordering of the prion-like domain of the Arabidopsis thermosensory protein EARLY FLOWERING 3. Proceedings of the National Academy of Sciences 120: e2304714120

Janitza P, Zhu Z, Anwer MA, van Zanten M, Delker C (2024) A Guide to Quantify Arabidopsis Seedling Thermomorphogenesis at Single Timepoints and by Interval Monitoring. Methods in Molecular Biology 2795: 3–16

Jones P, Binns D, Chang H-Y, Fraser M, Li W, McAnulla C, McWilliam H, Maslen J, Mitchell A, Nuka G (2014) InterProScan 5: genome-scale protein function classification. Bioinformatics 30: 1236–1240

Jung J-H, Barbosa AD, Hutin S, Kumita JR, Gao M, Derwort D, Silva CS, Lai X, Pierre E, Geng F (2020) A prion-like domain in ELF3 functions as a thermosensor in Arabidopsis. Nature 585: 256–260

Jung J-H, Domijan M, Klose C, Biswas S, Ezer D, Gao M, Khattak AK, Box MS, Charoensawan V, Cortijo S (2016) Phytochromes function as thermosensors in Arabidopsis. Science 354: 886–889

Kamioka M, Takao S, Suzuki T, Taki K, Higashiyama T, Kinoshita T, Nakamichi N (2016) Direct repression of evening genes by CIRCADIAN CLOCK-ASSOCIATED1 in the Arabidopsis circadian clock. The Plant Cell 28: 696–711

Kim W-Y, Fujiwara S, Suh S-S, Kim J, Kim Y, Han L, David K, Putterill J, Nam HG, Somers DE (2007) ZEITLUPE is a circadian photoreceptor stabilized by GIGANTEA in blue light. Nature 449: 356–360

Kumar SV, Lucyshyn D, Jaeger KE, Alós E, Alvey E, Harberd NP, Wigge PA (2012) Transcription factor PIF4 controls the thermosensory activation of flowering. Nature 484: 242–245

Laitinen RA, Nikoloski Z (2019) Genetic basis of plasticity in plants. Journal of Experimental Botany 70: 739–745

Lancaster AK, Nutter-Upham A, Lindquist S, King OD (2014) PLAAC: a web and command-line application to identify proteins with prion-like amino acid composition. Bioinformatics 30: 2501–2502

Larsson A (2014) AliView: a fast and lightweight alignment viewer and editor for large datasets. Bioinformatics 30: 3276–3278

Legris M, Klose C, Burgie ES, Rojas CCR, Neme M, Hiltbrunner A, Wigge PA, Schäfer E, Vierstra RD, Casal JJ (2016) Phytochrome B integrates light and temperature signals in Arabidopsis. Science 354: 897–900

Letunic I, Bork P (2007) Interactive Tree Of Life (iTOL): an online tool for phylogenetic tree display and annotation. Bioinformatics 23: 127–128

Lincoln C, Britton JH, Estelle M (1990) Growth and development of the axr1 mutants of Arabidopsis. The Plant Cell 2: 1071–1080

Linde AM, Eklund DM, Kubota A, Pederson ER, Holm K, Gyllenstrand N, Nishihama R, Cronberg N, Muranaka T, Oyama T (2017) Early evolution of the land plant circadian clock. New Phytologist 216: 576–590

Lindsay RJ, Wigge PA, Hanson SM (2023) Molecular basis of polyglutamine-modulated ELF3 phase change in Arabidopsis temperature response. bioRxiv: 2023.2003. 2015.532793

Liu XL, Covington MF, Fankhauser C, Chory J, Wagner DR (2001) ELF3 encodes a circadian clock–regulated nuclear protein that functions in an Arabidopsis PHYB signal transduction pathway. The Plant Cell 13: 1293–1304

Lu S, Zhao X, Hu Y, Liu S, Nan H, Li X, Fang C, Cao D, Shi X, Kong L (2017) Natural variation at the soybean J locus improves adaptation to the tropics and enhances yield. Nature genetics 49: 773–779

Lu SX, Webb CJ, Knowles SM, Kim SH, Wang Z, Tobin EM (2012) CCA1 and ELF3 Interact in the control of hypocotyl length and flowering time in Arabidopsis. Plant physiology 158: 1079–1088

Matasci N, Hung L-H, Yan Z, Carpenter EJ, Wickett NJ, Mirarab S, Nguyen N, Warnow T, Ayyampalayam S, Barker M (2014) Data access for the 1,000 Plants (1KP) project. Gigascience 3: 2047–2217X-2043-2017

Matsubara K, Ogiso-Tanaka E, Hori K, Ebana K, Ando T, Yano M (2012) Natural variation in Hd17, a homolog of Arabidopsis ELF3 that is involved in rice photoperiodic flowering. Plant and Cell Physiology 53: 709–716

McWatters HG, Bastow RM, Hall A, Millar AJ (2000) The ELF3 zeitnehmer regulates light signalling to the circadian clock. Nature 408: 716–720

Miller MA, Pfeiffer W, Schwartz T (2012) The CIPRES science gateway: enabling high-impact science for phylogenetics researchers with limited resources. In Proceedings of the 1st Conference of the Extreme Science and Engineering Discovery Environment: Bridging from the extreme to the campus and beyond, pp 1-8

Mizuno N, Matsunaka H, Yanaka M, Ishikawa G, Kobayashi F, Nakamura K (2023) Natural variations of wheat EARLY FLOWERING 3 highlight their contributions to local adaptation through fine-tuning of heading time. Theoretical and Applied Genetics 136: 139

Nakamichi N, Kiba T, Henriques R, Mizuno T, Chua N-H, Sakakibara H (2010) PSEUDO-RESPONSE REGULATORS 9, 7, and 5 are transcriptional repressors in the Arabidopsis circadian clock. The Plant Cell 22: 594–605

Nguyen L-T, Schmidt HA, Von Haeseler A, Minh BQ (2015) IQ-TREE: a fast and effective stochastic algorithm for estimating maximum-likelihood phylogenies. Molecular biology and evolution 32: 268–274

Nieto C, López-Salmerón V, Davière J-M, Prat S (2015) ELF3-PIF4 interaction regulates plant growth independently of the evening complex. Current Biology 25: 187–193

Nohales MA, Kay SA (2016) Molecular mechanisms at the core of the plant circadian oscillator. Nature structural & molecular biology 23: 1061–1069

Nusinow DA, Helfer A, Hamilton EE, King JJ, Imaizumi T, Schultz TF, Farré EM, Kay SA (2011) The ELF4– ELF3–LUX complex links the circadian clock to diurnal control of hypocotyl growth. Nature 475: 398–402

Park Y-J, Lee H-J, Gil K-E, Kim JY, Lee J-H, Lee H, Cho H-T, Vu LD, De Smet I, Park C-M (2019) Developmental programming of thermonastic leaf movement. Plant Physiology 180: 1185–1197

Press MO, Lanctot A, Queitsch C (2016) PIF4 and ELF3 act independently in Arabidopsis thaliana thermoresponsive flowering. PLoS One 11: e0161791

Prilusky J, Felder CE, Zeev-Ben-Mordehai T, Rydberg EH, Man O, Beckmann JS, Silman I, Sussman JL (2005) FoldIndex©: a simple tool to predict whether a given protein sequence is intrinsically unfolded. Bioinformatics 21: 3435–3438

Quint M, Delker C, Franklin KA, Wigge PA, Halliday KJ, Van Zanten M (2016) Molecular and genetic control of plant thermomorphogenesis. Nature plants 2: 1–9

Raschke A, Ibañez C, Ullrich KK, Anwer MU, Becker S, Glöckner A, Trenner J, Denk K, Saal B, Sun X (2015) Natural variants of ELF3 affect thermomorphogenesis by transcriptionally modulating PIF4-dependent auxin response genes. BMC plant biology 15: 1–10

Ridge S, Deokar A, Lee R, Daba K, Macknight RC, Weller JL, Tar’an B (2017) The chickpea Early Flowering 1 (Efl1) locus is an ortholog of Arabidopsis ELF3. Plant physiology 175: 802–815

Ronald J, Davis SJ (2019) Focusing on the nuclear and subnuclear dynamics of light and circadian signalling. Plant, Cell & Environment 42: 2871–2884

Ronald J, Su C, Wang L, Davis SJ (2022) Cellular localization of Arabidopsis EARLY FLOWERING3 is responsive to light quality. Plant Physiology 190: 1024–1036

Ronald J, Wilkinson AJ, Davis SJ (2021) EARLY FLOWERING3 sub-nuclear localization responds to changes in ambient temperature. Plant Physiology 187: 2352–2355

Rozas J, Ferrer-Mata A, Sánchez-DelBarrio JC, Guirao-Rico S, Librado P, Ramos-Onsins SE, Sánchez-Gracia A (2017) DnaSP 6: DNA sequence polymorphism analysis of large data sets. Molecular biology and evolution 34: 3299–3302

Sabate R, Rousseau F, Schymkowitz J, Ventura S (2015) What makes a protein sequence a prion? PLoS computational biology 11: e1004013

Saito H, Ogiso-Tanaka E, Okumoto Y, Yoshitake Y, Izumi H, Yokoo T, Matsubara K, Hori K, Yano M, Inoue H (2012) Ef7 encodes an ELF3-like protein and promotes rice flowering by negatively regulating the floral repressor gene Ghd7 under both short-and long-day conditions. Plant and Cell Physiology 53: 717–728

Tajima F (1989) Statistical method for testing the neutral mutation hypothesis by DNA polymorphism. Genetics 123: 585–595

Tajima T, Oda A, Nakagawa M, Kamada H, Mizoguchi T (2007) Natural variation of polyglutamine repeats of a circadian clock gene ELF3 in Arabidopsis. Plant biotechnology 24: 237–240

Toombs JA, McCarty BR, Ross ED (2010) Compositional determinants of prion formation in yeast. Molecular and cellular biology 30: 319–332

Toombs JA, Petri M, Paul KR, Kan GY, Ben-Hur A, Ross ED (2012) De novo design of synthetic prion domains. Proceedings of the National Academy of Sciences 109: 6519–6524

Undurraga SF, Press MO, Legendre M, Bujdoso N, Bale J, Wang H, Davis SJ, Verstrepen KJ, Queitsch C (2012) Background-dependent effects of polyglutamine variation in the Arabidopsis thaliana gene ELF3. Proceedings of the National Academy of Sciences 109: 19363–19367

Uversky VN (2019) Supramolecular fuzziness of intracellular liquid droplets: liquid–liquid phase transitions, membrane-less organelles, and intrinsic disorder. Molecules 24: 3265

van Zanten M, Voesenek LA, Peeters AJ, Millenaar FF (2009) Hormone-and light-mediated regulation of heat-induced differential petiole growth in Arabidopsis. Plant Physiology 151: 1446–1458

Walters RH, Murphy RM (2009) Examining polyglutamine peptide length: a connection between collapsed conformations and increased aggregation. Journal of molecular biology 393: 978–992

Wang L, Fujiwara S, Somers DE (2010) PRR5 regulates phosphorylation, nuclear import and subnuclear localization of TOC1 in the Arabidopsis circadian clock. The EMBO journal 29: 1903–1915

Wilkinson EG, Strader LC (2020) A Prion-based thermosensor in plants. Molecular Cell 80: 181–182

Wittern L, Steed G, Taylor LJ, Ramirez DC, Pingarron-Cardenas G, Gardner K, Greenland A, Hannah MA, Webb AA (2023) Wheat EARLY FLOWERING 3 affects heading date without disrupting circadian oscillations. Plant Physiology 191: 1383–1403

Xu X, Zheng C, Lu D, Song CP, Zhang L (2021) Phase separation in plants: new insights into cellular compartmentalization. Journal of Integrative Plant Biology 63: 1835–1855

Yamaguchi R, Nakamura M, Mochizuki N, Kay SA, Nagatani A (1999) Light-dependent translocation of a phytochrome B-GFP fusion protein to the nucleus in transgenic Arabidopsis. The Journal of cell biology 145: 437–445

Zahn T, Zhu Z, Ritoff N, Krapf J, Junker A, Altmann T, Schmutzer T, Tüting C, Kastritis PL, Babben S (2023) Novel exotic alleles of EARLY FLOWERING 3 determine plant development in barley. Journal of Experimental Botany: erad127

Zakhrabekova S, Gough SP, Braumann I, Müller AH, Lundqvist J, Ahmann K, Dockter C, Matyszczak I, Kurowska M, Druka A (2012) Induced mutations in circadian clock regulator Mat-a facilitated short-season adaptation and range extension in cultivated barley. Proceedings of the National Academy of Sciences 109: 4326–4331

Zambrano R, Conchillo-Sole O, Iglesias V, Illa R, Rousseau F, Schymkowitz J, Sabate R, Daura X, Ventura S (2015) PrionW: a server to identify proteins containing glutamine/asparagine rich prion-like domains and their amyloid cores. Nucleic acids research 43: W331–W337

Zhou L, Feng T, Xu S, Gao F, Lam TT, Wang Q, Wu T, Huang H, Zhan L, Li L (2022) ggmsa: a visual exploration tool for multiple sequence alignment and associated data. Briefings in Bioinformatics 23: bbac222

Zhu Z, Esche F, Babben S, Trenner J, Serfling A, Pillen K, Maurer A, Quint M (2023) An exotic allele of barley EARLY FLOWERING 3 contributes to developmental plasticity at elevated temperatures. Journal of Experimental Botany 74: 2912–2931

Zhu Z, Quint M, Anwer MU (2022) Arabidopsis EARLY FLOWERING 3 controls temperature responsiveness of the circadian clock independently of the evening complex. Journal of Experimental Botany 73: 1049–1061

